# 4D Cell-Condensate Bioprinting

**DOI:** 10.1101/2022.02.28.482216

**Authors:** Aixiang Ding, Rui Tang, Felicia He, Sang Jin Lee, Kaelyn Gasvoda, Eben Alsberg

## Abstract

4D bioprinting techniques that facilitate formation of shape-changing scaffold-free cell condensates with prescribed geometries have yet been demonstrated. Here, a simple yet novel 4D bioprinting approach is presented that enables formation of a shape-morphing cell condensate-laden bilayer system comprised of an actuation layer and a cell condensate-supporting microgel (MG) layer. The strategy produces scaffold-free cell condensates which morph over time into predefined complex shapes. With a sequential printing (i.e., MG printing first onto the preformed actuation hydrogel layer and cell-only printing inside the pre-printed MG construct second), cell condensate-laden bilayers with specific geometries are readily fabricated and can be further UV-crosslinked to form strong interlayer adhesion. Since the bilayers have tunable deformability and MG degradation can be tailored, this enables controllable morphological transformations and on-demand liberation of cell condensates. With this system, large cell condensate-laden constructs with various complex shapes were obtained through predefined conformational conversions. As a proof-of-concept study, the formation of the letter “C” and helix-shaped robust cartilage-like tissues differentiated from human mesenchymal stem cells (hMSCs) was demonstrated. This new system brings about a new versatile 4D bioprinting platform idea that is anticipated to broaden and facilitate the applications of cell condensation-based 4D bioprinting.

## 1. Introduction

Since it was first introduced by Tibbits from the Massachusetts Institute of Technology in 2013,^[1]^ four-dimensional (4D) printing,^[2]^ as a newly emerging technology platform built on the well-known three-dimensional (3D) printing (also referred to as additive manufacturing, AM),^[3]^ has received rapidly increasing research interest in the last few years due to its enormous application prospects in a myriad of fields including soft robotics,^[4]^ actuators,^[5]^ flexible wearable electronics,^[6]^ and biomedical engineering.^[7]^ Similar to 3D printing, 4D printing relies on the selection of materials, printer, and mathematical modeling and design.^[8]^ However, since 4D printing adopts specifically smart materials that respond to external stimuli (e.g., light,^[9]^ moisture,^[10]^ temperature,^[11]^ electric fields,^[12]^ and magnetic fields^[13]^) through a change in the macroscopic morphology,^[14]^ an additional time dimension is integrated with the 3D printing to endow the printed architecture with the ability to undergo spatiotemporal shape transformation,^[15]^ affording constructs with more sophisticated configurations,^[16]^ and more importantly, enabling the capability to reshape over time in accordance with practical needs.^[17]^ As such, 4D printing shows a clear advantage over 3D printing for tissue regeneration-related applications, where time-dependence is an important consideration for recapitulating the structural evolution of conformationally intricate tissues during maturation.^[18]^

4D bioprinting that involves the use of biocompatible materials and incorporation of viable cells^[19]^ specializes in the fabrication of bioactive cell-laden constructs (bioconstructs) capable of dynamic reconfiguration.^[20]^ The smart materials, material processing, and stimulation must be strictly cytocompatible to maintain viability of incorporated cells.^[21]^ Even though a variety of materials such as alloys,^[22]^ ceramics,^[23]^ liquid crystal elastomers,^[24]^ composites,^[25]^ and shape-memory polymers^[26]^ have been widely utilized to date in 4D printing, they generally fail to meet the demands of live-cell encapsulation.^[27]^ In this regard, given that hydrogel scaffolds can provide a soft tissue-like environment to support cell growth,^[28]^ efforts have been dedicated to developing hydrogel-based materials that allow for shape change under physiological stimulation for 4D bioprinting.^[29]^ For example, hollow self-folding tubes were fabricated using alginate and hyaluronic acid-derived hydrogels,^[30]^ and gelatin and polyethylene glycol-based hydrogels were employed to fabricate a 4D physiologically adaptable patch to treat myocardial infarction.^[31]^ An alginate/polydopamine composite was used as cell-laden bioink to fabricate near-infrared-triggered shape-morphing constructs.^[32]^ In a recent study, 4D digital light printed (DLP) silk hydrogels were applied as implants for treatment of a damaged trachea.^[33]^ We also developed a series of biopolymer-based^[34]^ and microgel-based^[35]^ bioinks as versatile bioprinting platforms.

Despite inspiring preliminary achievements in 4D bioprinted cell-laden hydrogels, the scaffold-based approach often encounters the difficulty of synchronizing hydrogel degradation with the new tissue formation,^[36]^ presents potential cytotoxicity originating from the fabrication techniques and degradation byproducts,^[37]^ and strongly restricts the cell-to-cell interactions that are important during tissue development and healing.^[38]^ In comparison, scaffold-free, cell condensation-based alternatives exclude the use of scaffolding materials for individual cell encapsulation to yield a tissue construct that can readily circumvent the above limitations.^[39]^ Taking the importance of shape transformations and remodeling during tissue formation into consideration, the development of a cell condensation construct that bears programmable deformation capability after fabrication is of great significance. Nonetheless, a system with this capability has not been demonstrated yet.

To address this challenge, a 4D cell-condensate bioprinting strategy using a unique bilayered system has been developed in this work to impart the shape-morphing feature to a 3D printed cell construct (**Scheme 1**). The proposed bilayer consists of a preformed gradient-crosslinking hydrogel layer as the actuation layer that drives the shape morphing and a printed photocurable and degradable cell-supporting microgel (MG) layer that allows printing a cell-only bioink inside and maintains the shape of the printed cells as they form a condensate in the initial stage. Upon the gradual and controllable degradation of the photocrosslinked MG layer during culture in tissue specific differentiation media, 4D transformation into a mature tissue with a well-defined configuration can be obtained on demand.

**Scheme 1.**
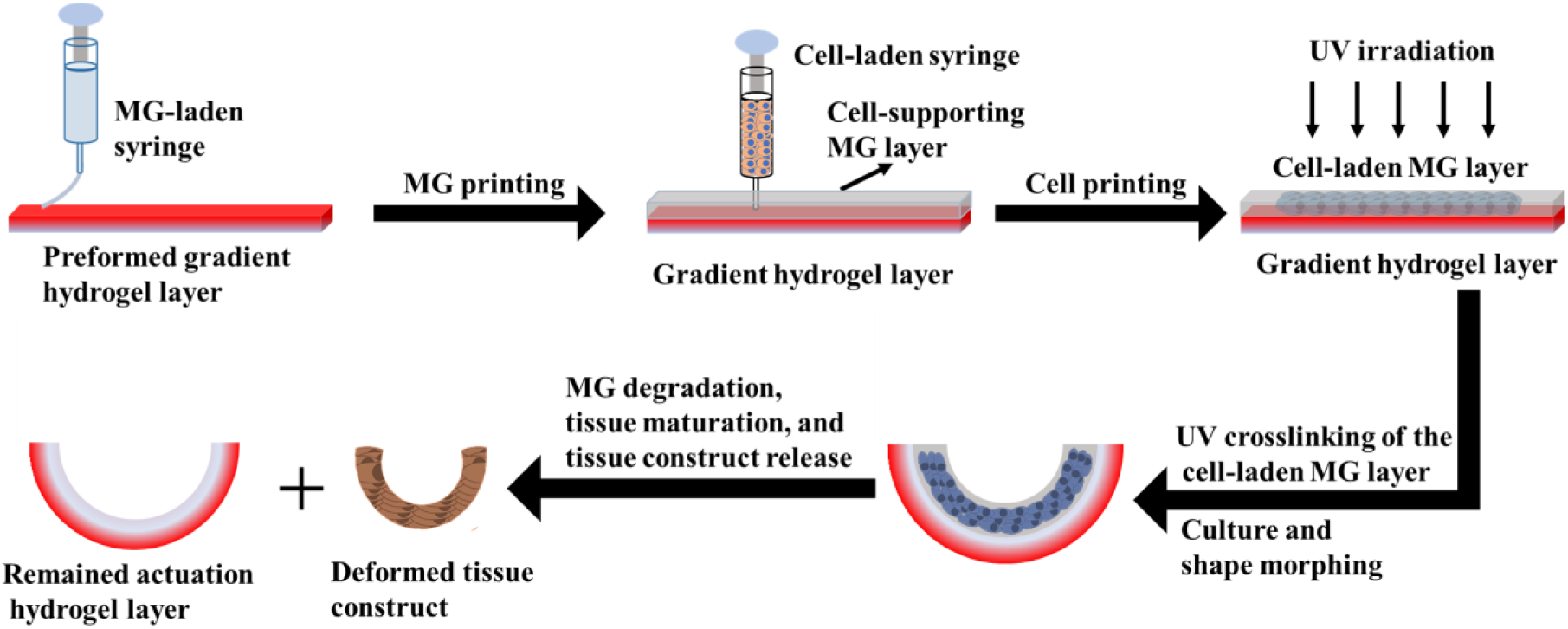
Schematic illustration of 4D cell-condensate bioprinting, tissue maturation, and tissue release.

## 2. Results and discussion

### 2.1. Cell-free bilayer fabrication and shape-morphing behaviors

We have previously demonstrated the feasibility of incorporation of a UV absorber into a single-component photocrosslinkable biopolymer, such as an oxidized and methacrylated alginate (OMA, Figure S1a), to effectively generate a crosslinking gradient throughout the hydrogel thickness by a one-step UV photocrosslinking (Figure S1c) (supporting information),^[34]^ which is attributed to UV absorber-based light attenuation along the light pathway.^[40]^ Consequently, an internal anisotropic strain was generated after swelling in media, leading to hydrogel deformation (Figure S1c). Herein, a bicomponent photocrosslinkable biopolymer composite of O5M45A (OMA with 5% theoretical oxidation and 45% theoretical methacrylation) and GelMA (methacrylated gelatin) (Figure S1b) was employed to form a gradient hydrogel [O5M45A/GelMA(g)]. While the bicomponent hydrogel without gradient formation (O5M45A/GelMA) exhibited a similar modulus with the single-component non-gradient hydrogel (O5M45A), the elastic modulus of O5M45A/GelMA(g) was smaller than that of O5M45A(g) (**Figure 1a**), which implies that O5M45A/GelMA(g) had a larger gradient range in comparison with the single-component gradient hydrogel [O5M45A(g)]. As a result, this bicomponent gradient hydrogel exhibited greater deformability (Figure S2). The printability of O5M20A microgels (MGs), which are calcium ion (Ca^2+^)-crosslinked hydrogels fabricated according to our previously described method (supporting information),^[41]^ was then studied prior to the bilayer hydrogel fabrication. The prepared MGs behaved as a stable bulk hydrogel under low shear strains, (Figure 1b and S3a) but yielded at a shear strain over 25% (Figure 1b). Shear-thinning behavior was also identified by increasing the shear rate (Figure 1c), shear strain (Figure S3b), and shear stress (Figure S3c). Moreover, the MGs displayed a rapid and reversible phase transition between the solid-like (elastic) state and the liquid-like (viscous) state by alternating the shear strain applied between 1% and 100% (Figure 1d and S3d), suggesting favorable extrudability and rapid self-healing after deposition.^[41, 42]^ Consequently, MG structures with high resolution were readily printed (Figure S4a) and further stabilized by subsequent UV-crosslinking (Figure S4b), resulting in dual-crosslinked constructs. As expected, due to the low methacrylation degree, the MGs exhibited a rapid and tunable degradation profile in cell growth media. For example, the dual-crosslinked MG completely degraded in 14 days when UV-crosslinked for 20 s, while it took approximately 28 days for complete degradation if the MGs were UV irradiated for 30 s (Figure 1e). This result provides evidence that the liberation of a cell condensation construct at a predetermined time point may be accomplished by simply adjusting the UV-crosslinking time. In addition, the UV-crosslinked MGs also exhibited a higher swelling ratio (Figure 1f) but much weaker mechanics (Figure 1a) than the non-MG gradient hydrogels at the initial stage, and the swelling and mechanics decreased along with the degradation over time (Figure 1f and S5). In contrast, the O5M45A/GelMA(g) was relatively stable in the media during the course of a 28-day culture. The higher stability of the proposed O5M45A/GelMA(g) actuation layer can provide a stable shape-morphing force to maintain the shape of the deformed cell construct for long-term culture.

**Figure 1.**
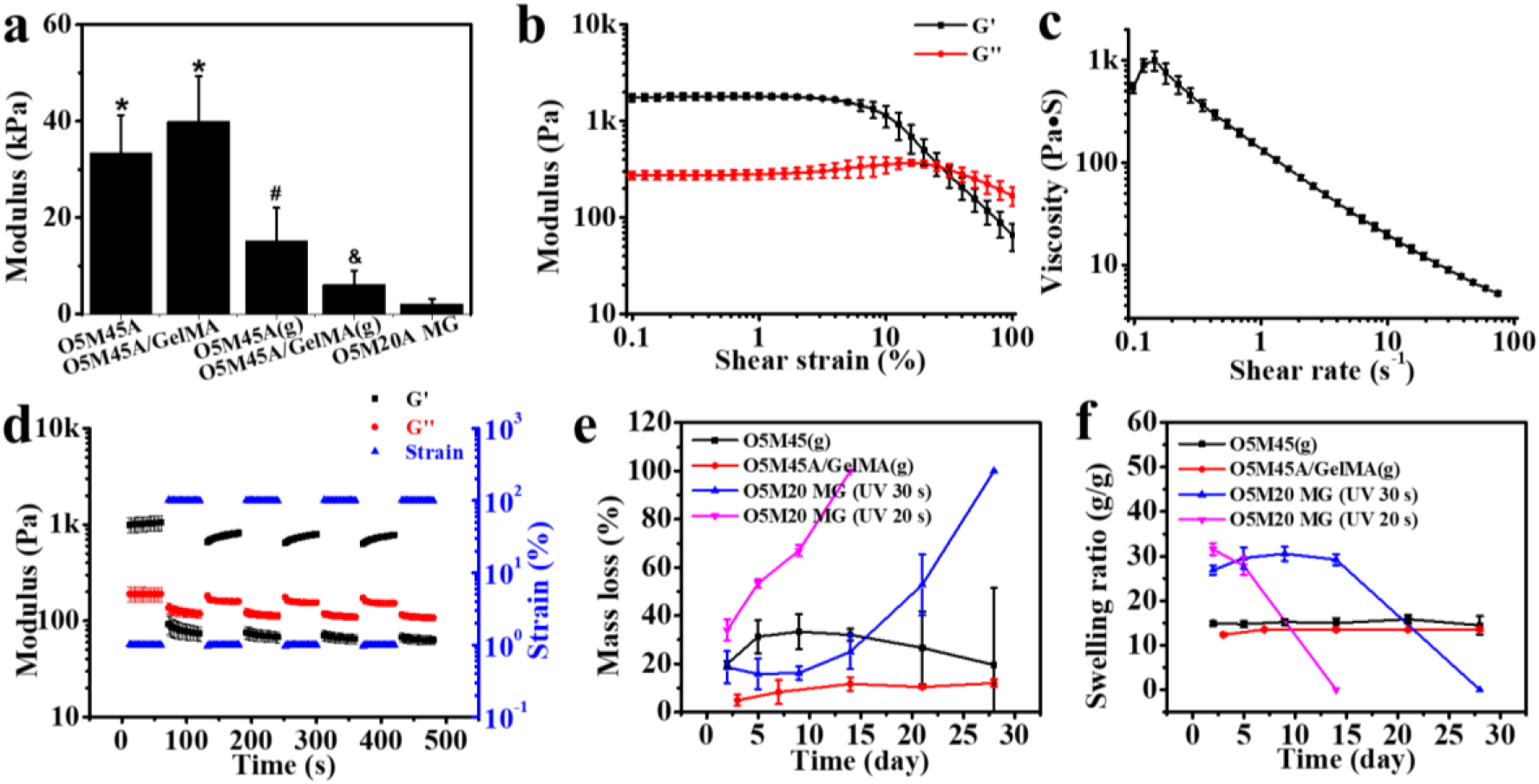
(a) Compressive elastic moduli of various hydrogels, and the change of (b) modulus and (c) viscosity of O5M20A MGs with increasing shear strain and shear rate, respectively. (d) The rapid and reversible phase transition of O5M20A MGs between elastic state and viscous state by alternating the shear strain between 1% and 100%, and the (e) degradation and (f) swelling profiles of various hydrogels. *^, #, &^*p* < 0.05 compared to group sharing no or a different symbol.

Owing to its printability and photocrosslinkability, the O5M20A MGs can be finely printed using a 22-gauge needle onto the surface of a pre-fabricated O5M45A/GelMA(g) substrate (Movie S1). The printed MG layer was subsequently UV-crosslinked, forming a stable bilayer system with robust interfacial adhesion between the two layers (**Figure 2a, 2b**, and S6) through the covalent crosslinking of the remaining methacrylate groups on the preformed O5M45A/GelMA(g) surface (Figure S7) with the methacrylate groups on the O5M20A MG.^[43]^ Although the as-crosslinked MGs exhibited a higher swelling ratio than the O5M45A/GelMA(g) hydrogel (Figure 1f), this bilayer hydrogel (i.e., a disc) rapidly rolled into a tubular structure towards the MG side when immersed in the phosphate buffered saline (PBS, pH 7.4) media (Figure 2c and Movie S2), due to a synergistic contribution of (i) the large gradient within the actuation layer that generates a substantial actuation force large enough to overcome the opposite layer strain brought by the swelling discrepancy and (ii) the extremely weak stiffness of the MG layer that allows easy deformation. The role of the actuation layer in driving the shape-morphing process was further verified by a comparison experiment (Figure S8). Among hydrogel strips tested, including single-layer non-gradient O5M45A/GelMA, gradient O5M45A/GelMA and MG hydrogels (Figure S8a−c) and bilayer hydrogels (Figure S8d−e) with a non-gradient or gradient O5M45A/GelMA layer, only the hydrogels with a gradient layer showed remarkable rolling, while the shapes of the hydrogels involving no gradient remained unchanged. Interestingly, the bilayer that consisted of an MG layer and a non-gradient O5M45A/GelMA (Figure S8d) did not show any shape change regardless of the swelling difference between the two layers. This is because the MG layer is too soft to serve as either an actuation layer or a shape-constraint layer. For this reason, the bilayer composed of an MG layer and a gradient O5M45A/GelMA layer shared a comparable bending angle with the single-layer gradient hydrogel (Figure S8f). This exquisite design offers a unique bilayer system in which the two layers have specific, independent functions.

**Figure 2.**
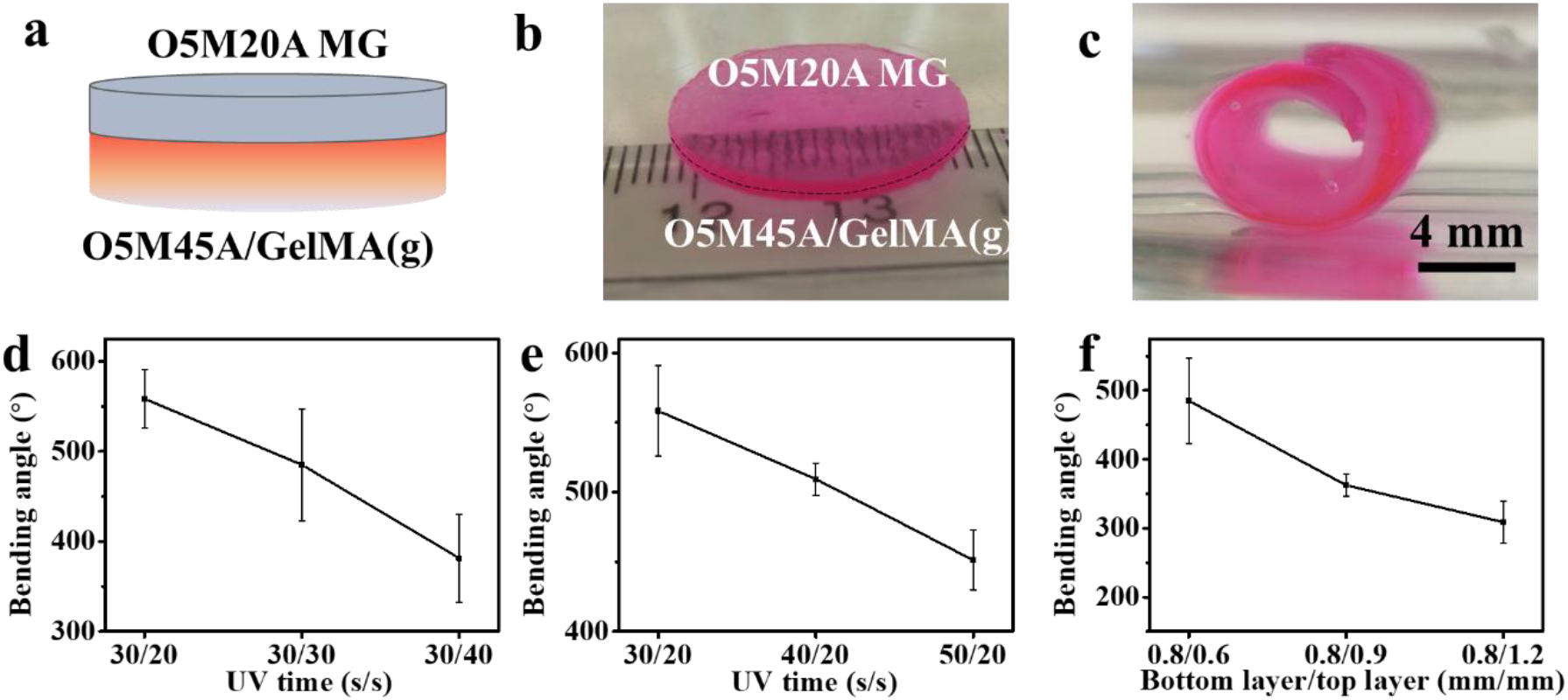
(a) Illustration and real sample photographs of a bilayer hydrogel disc (b) before and (c) after shape morphing. Effects of varying UV time applied to (d) the upper layer and (e) the bottom layer and (f) of varying MG layer thickness on the bending behaviors of the hydrogel strips. The hydrogel strip thickness for the UV time variation study was set to 0.8/0.5 (mm/mm) and the UV time for the MG layer thickness variation study was set to 30s/30s.

The prototype of the bilayer hydrogel strip was then used to investigate the impact of UV crosslinking time exposed to each layer (the gradient layer and the MG layer) and the thickness of the MG layer on shape-morphing behavior. Results of the quantified bending angles revealed a clear influence of these parameters on the hydrogel deformation (Figure 2d∼e). Increasing UV exposure time and MG layer thickness both exerted negative effects on the hydrogel bending. As far as the UV time, the receiving time for each layer can be selectively tuned to control the hydrogel bending. For example, the UV time for the upper layer (MG layer) can be adjust after being deposited onto the preformed gradient layer (Figure 2d and S9a) or the UV time can be adjusted when fabricating the bottom layer (gradient layer) while fixing the UV time for the upper layer (Figure 2e and S9b). The lowered bending arose from the combined outcome of reduced gradient range and lowered volumetric swelling by increasing UV time.^[33]^ Differing from the UV time, the lowered bending achieved by the increasing MG thickness (Figure 2f and S9c) can be mainly attributed to increased resistance of the MG layer to the morphing. This controllable shape-morphing property will benefit applications where the capacity to tailor the extent of geometric change is desirable.

### 2.2 Cell condensate-laden bilayer fabrication and their shape-morphing behaviors

The printability of the live cells within the MG layer was examined before fabricating the cell condensate-laden bilayer system. Live cells themselves can serve as a cell-only bioink^[41]^ to be printed into an O5M20A supporting MG bath, which was first printed as described above. In this cell-only printing process, the supporting material is typically shear-thinning and rapidly self-healing,^[41, 42]^ permitting free embedding and deposition of live cells by replacing the MGs along the needle moving pathway and concurrently maintaining the printed cell construct with high fidelity. After printing, the dual-crosslinkable property of the MGs enables UV crosslinking to further stabilize the printed cell-only bioink and permits cell condensation formation without an intervening scaffold material. HeLa cells were first printed into a cell filament with a 25-gauge needle inside an as-printed MG strip (22 × 5 × 1 mm^3^). The cell filament-laden MG strip was subsequently UV crosslinked and imaged under a microscope to examine the printing resolution and then cultured in cell-growth media for 4 h to examine the cell viability. Results showed that the cells were printed into a filament with high resolution (Figure S10), confirming reliable cell printability, and remained highly viable (**Figure 3a**), suggesting no obvious adverse effects of the bioprinting process and UV crosslinking on cell survival. Next, larger cell constructs, e.g., a cell strip (18 × 4 × 1.2 mm^3^, Figure 3b,left and middle images), a cell sheet (13 × 13 × 1.2 mm^3^, Figure 3b,right image, and Figure S11), and a cell cuboid (5 × 5 × 4 mm^3^, Figure S12), with high resolution were then printed into the supporting MG bath, further confirming the cell printability.

**Figure 3.**
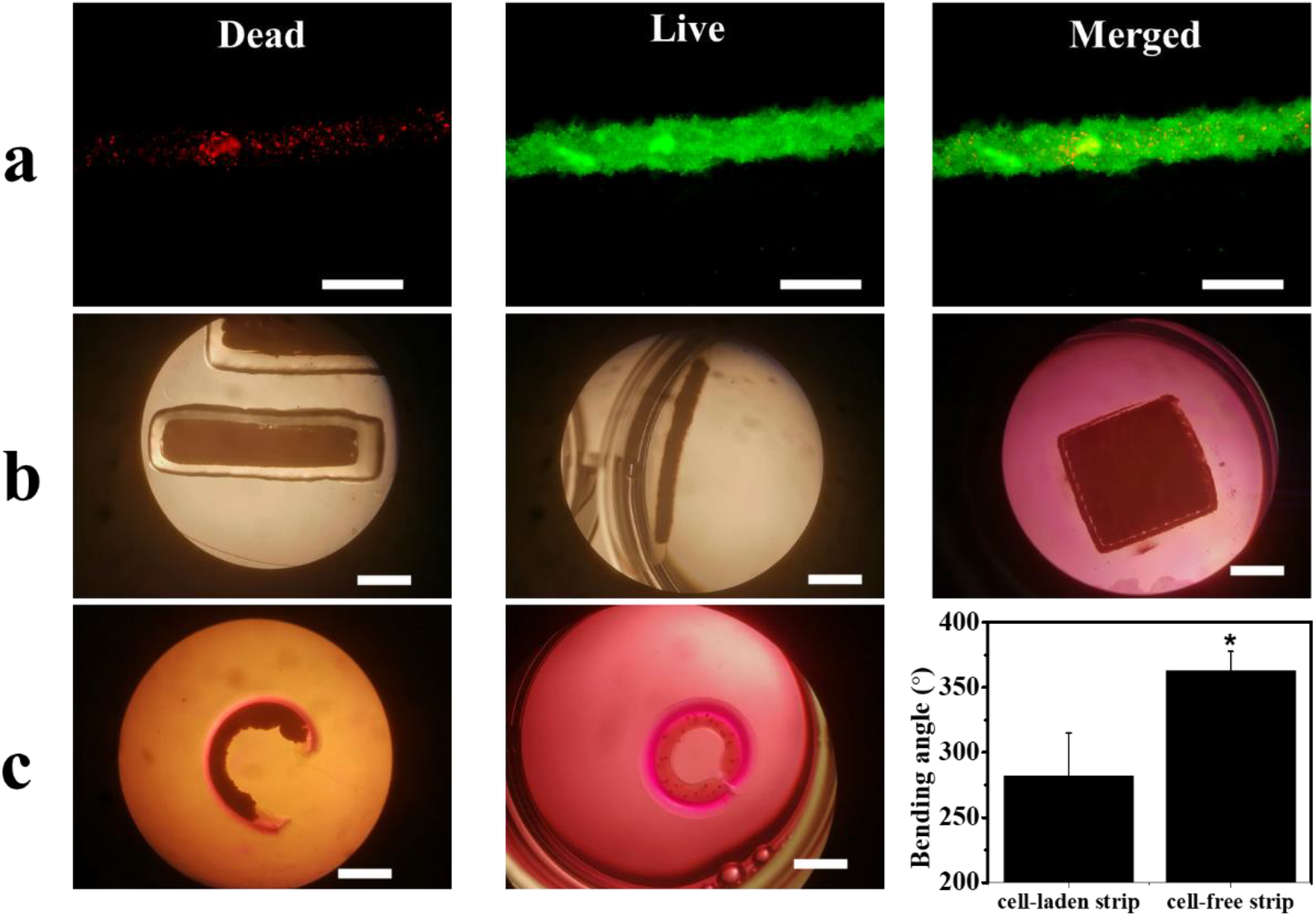
(a) Live/dead cell staining of a printed cell filament inside photocrosslinked MGs after culturing in cell growth media for 4 h. Scale bars indicate 0.5 mm. (b) Photomicrographs of a printed cell strip (18 × 4 × 1.2 mm^3^) from top view (left) and side view (middle) and a cell sheet (13 × 13 × 1.2 mm^3^, right) inside photocrosslinked MG constructs. The cell strip images were taken immediately after UV photocrosslinking, the cell sheet image was taken immediately after placing in cell growth media. Scale bars indicate 5 mm. (c) Representative photomicrographs of cell-laden (left) and cell-free (middle) bilayer strips after culturing in cell growth media for 4 h, and quantified bending angles of the cell-laden and cell-free bilayer strips (right). \**p* < 0.05 and scale bars indicate 5 mm. The UV irradiation times for the gradient layer and the MG layer fabrications were set to 30 s and 20 s (30s/20s), respectively.

Cell condensate-laden bilayer hydrogel strips were then fabricated by a sequential printing process (30s/20s UV crosslinking time) and the shape-morphing behaviors were recorded. The O5M20A MG strip (22 × 5 × 1.2 mm^3^) was first printed onto the surface of a preformed O5M45A/GelMA(g) hydrogel disc and cells were subsequently printed within the MG layer to form a cell strip-laden bilayer construct (Movie S3), which was then UV crosslinked. Cell condensate-laden bilayer constructs were then obtained by cutting from the large construct and cultured in cell growth media for 4 h until no further morphological changes could be observed. As expected, the cell-laden bilayer strip underwent a bending process into a letter “C” shape towards the cell-laden layer side (Figure 3c,left) with, however, a smaller bending angle in comparison to the cell-free bilayer strip (Figure 3c,middle and right), most likely due to enhanced morphing resistance by the infilled cell condensate. Noteworthily, the shape-morphing process did not compromise the integrity of the printed cell construct because of the firm support by the surrounding dual-crosslinked MGs.

### 2.3 Printing of large cell constructs with complex structures and their shape-morphing behaviors

The successful 4D bioprinting of the bilayer cell condensate-laden hydrogel strips inspired printing of larger cell condensate-laden constructs with more complex structures and investigation of their shape-morphing behaviors. Four different bilayer bioconstructs, including sheet-based bilayers with a cell filling in the shape of a sheet, bar grid, and net, and a disc-based bilayer with a disc-shaped cell infilling, were fabricated with a similar sequential printing process described above (**Figure 4**, 40s/20s UV crosslinking time). HeLa cells were printed into specific geometries within the MG layer with high resolution, and these self-transformable cell condensate-laden and cell-free constructs morphed into concave structures by curling up to the cell layer side after culturing in cell growth media for 4 h. In addition, these bioconstructs appeared to be less curled compared to their cell-free counterparts, in agreement with the results obtained from the bilayer hydrogel strips. The above results collectively demonstrated the feasibility and effectiveness of this strategy to fabricate cell condensates with prescribed configurations by the 4D bioprinting technique.

**Figure 4.**
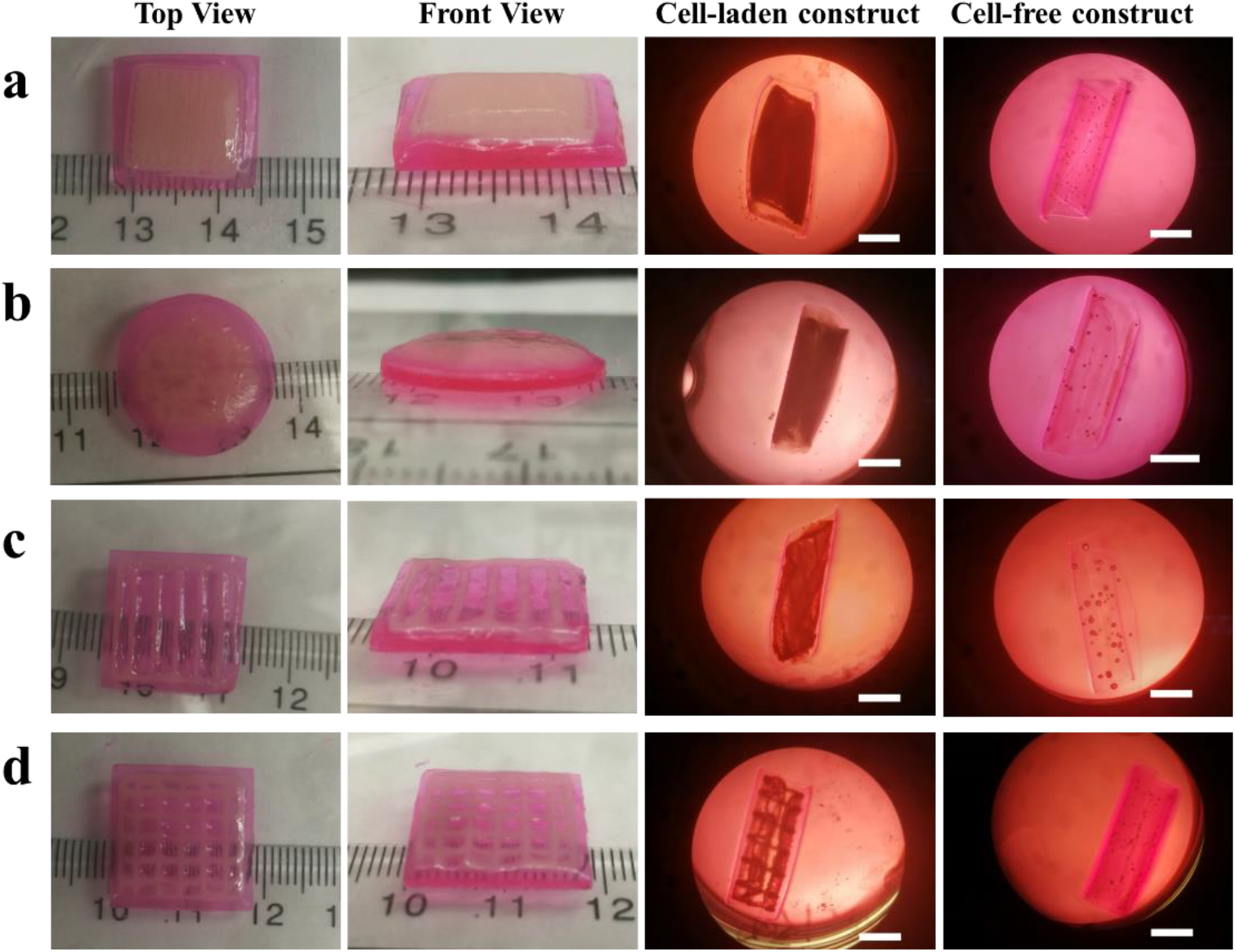
Large cell-laden bilayer constructs with defined structures of cell filling and the corresponding deformed configurations after culturing in cell growth media for 4 h: (a) sheet, (b) disc, (c) bar grid, and (d) net. Deformed cell-free bilayer counterparts were obtained under the same conditions and included for comparison. HeLa cells were used in this study, and UV irradiation times for the actuation layer (bottom layer) and the cell-laden layer (upper layer) were set to 40 s and 20 s (40s/20s), respectively. Scale bars indicate 5 mm.

To assess the long-term cell viability and stability of the bioconstructs, the cell-net infilled bilayer sheet (Figure 4d) was cultured in cell growth media, cell viability was examined using a live/dead staining assay and shape changes were monitored over a course of 14 days (Figure 5). The curling of the bioconstruct progressively increased during the first few days of culture (before day 5, Figure S13) due to MG layer degradation-induced softening of the cell-laden layer (Figure S5). The overall structure remained in a cylindrical shape, and the printed cells went through condensation to reinforce the “cell-net filament” during this period. It is interesting to note that the two layers (*i*.*e*., the actuation layer and cell condensate-laden layers) were observed to have visibly separation after a media change on day 6 while the structures of the individual layers were maintained (**Figure 5a** and S14a, D6). The layer separation stemmed from the degradation of the interfacial covalent crosslinks between the two layers. However, the shape of the cell condensate-laden MG layer was still stable because most crosslinked MGs were still retained at this time point. The gradually increased degradation of the MG layer with increasing culture time ultimately resulted in the disintegration of the MG layer on day 9 and some of the cell-net filaments were “liberated” from the MG layer (Figure 5a,D9). After continued culture of the bioconstruct to day 12, the MG layer was found to have further disintegrated, (Figure 5a,D12) and it was difficult for the bioconstruct to withstand the media change-mediated disturbance. With further culturing, the dual-crosslinked MGs almost completely degraded and were unable to support the cell-net structure, which eventually crumbled on day 14 (Figure S14b, D14). During the entire culturing period, the cell-net filaments maintained integrity, regardless of the MG degradation (Figure 5c∼5d, bright-field images), suggesting that strong physical cell-cell cadherin interactions were present within the cell condensate, and the cells were highly viable (Figure 5c∼5d, live/dead stained images), indicative of good cytocompatibility of this system. As the MG degradation can be tuned by controlling UV crosslinking time, in another independent experiment, a cell-net infilled bilayer construct was fabricated by increasing the UV exposure time for the MG layer from 20 s to 30 s, and the obtained bioconstruct (40s/30s UV crosslinking time) went through a similar but slower 4D process to form a tubular structure and maintained stable configuration over 21 days (Figure S15).

**Figure 5.**
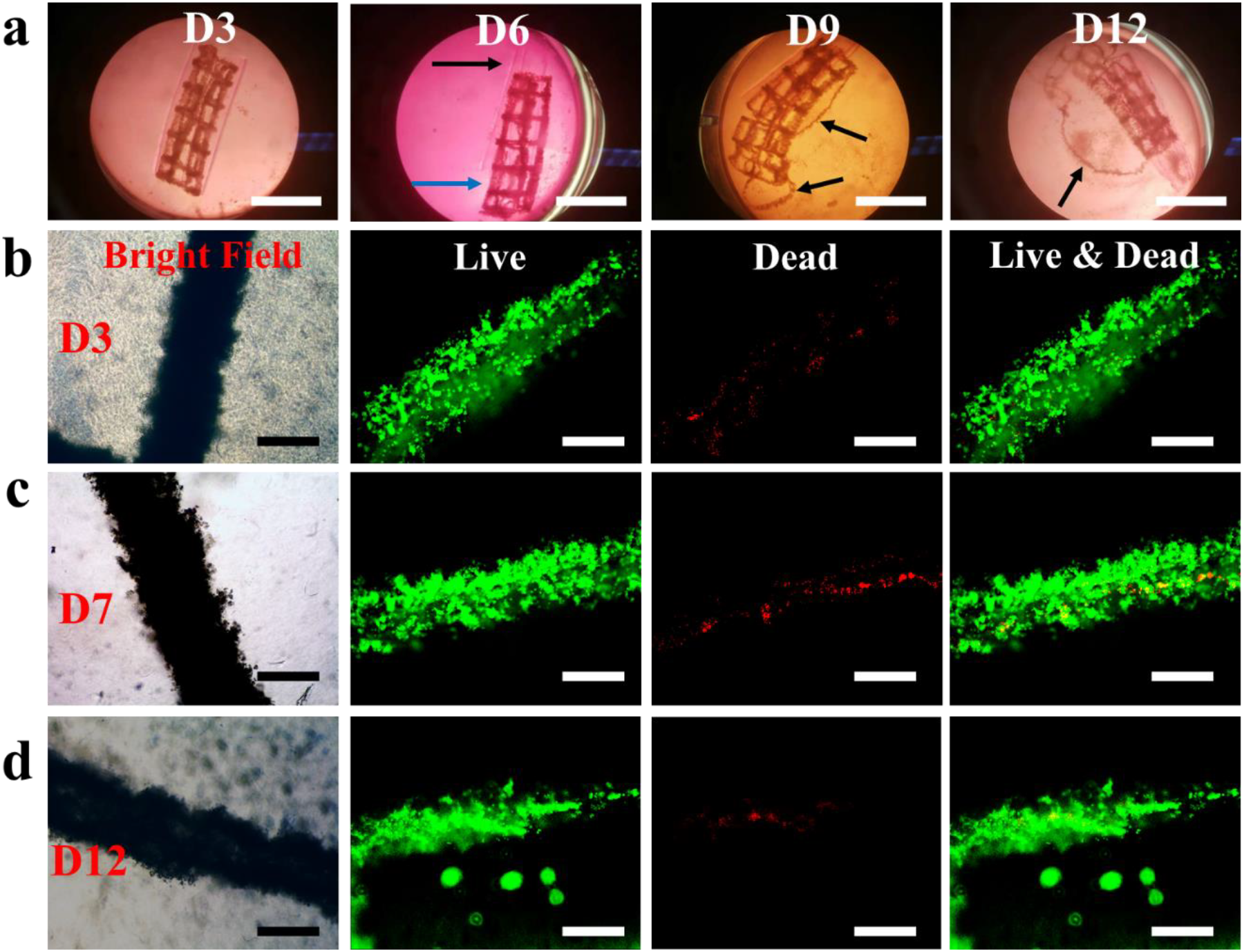
(a) Photomicrographs of cell-net infilled bilayer sheet at D3, D6, D9, and D12. The black arrow and blue arrow in the image at D6 show the separated gradient hydrogel layer and the cell condensate-laden layer, respectively. Arrows in images of D9 and D12 show “liberated” cell-net filaments. Scale bars indicate 10 mm. Photomicrographs showing a cell-net filament inside the hydrogel under bright-field channel, green (live) channel, red (dead) channel, and merged channel of green and red at (b) D3, (c) D7, and (d) D12. Scale bars indicate 0.5 mm. UV crosslinking time was set to bottom/top 40s/20s.

### 2.4 4D ex vivo engineering cartilage-like tissue with pre-programmed shape

With this system, specific tissues with well-defined configurations through a reshaping process can be obtained after culturing the cell-condensate construct in a cell-differentiation environment for an appropriate time course. To explore the application potential of this 4D-condensate bioprinting system in tissue engineering, a proof-of-concept study was performed by fabricating hMSC cell condensate-laden bilayer strips (50 s/20 s UV crosslinking time) with this 4D bioprinting strategy and then culturing the bilayer strips in chondrogenic media for 21 days to generate hydrogel-free 4D tissues. Like the progressive shape-morphing behaviors observed in the aforementioned sheet-shape-based systems, the bilayer strips in the experimental groups (Exp) deformed into a letter “*C*” shape in the first 24 h culture and the bending slowly increased until 14 days. Then the structures remained stable from 14 to 21 days (**Figure 6a, 6c**, and S16). Since the two layers detached at D7, we speculate the limited increased bending after D7 may have resulted from cell cytoskeleton-based contractile forces within the cell condensates. This similar phenomenon was also observed in the controls (cell condensate-laden single-layer MG strips, Ctrl), although minimal bending occurred in this group (Figure 6a, 6c, and S17). Cartilage-like tissues in the shapes of the Letter “*C*” (Figure 6b) and a nearly straight line (Figure S18) were finally obtained after full degradation of the supporting MGs in the Exp and Ctrl groups, respectively, after culture for 21 days (Figure S19). The 4D engineered cartilage-like tissues presented a similar level of glycosaminoglycan (GAG) production (Figure 6d), a key cartilage extracellular matrix component, and similar Young’s modulus compared to the cartilage-like tissue obtained from the controls (Figure 6e). Hematoxylin and eosin (H&E) staining showed that the engineered constructs exhibited a homogeneous pattern of tissue comprised of uniformly distributed chondrocytes. Strong safranin O (SafO) and toluidine blue O (TBO) staining in the 4D constructs also revealed substantial GAG production (Figure 6f∼6h) and appeared similar to the staining in the control (Figure S20).

**Figure 6.**
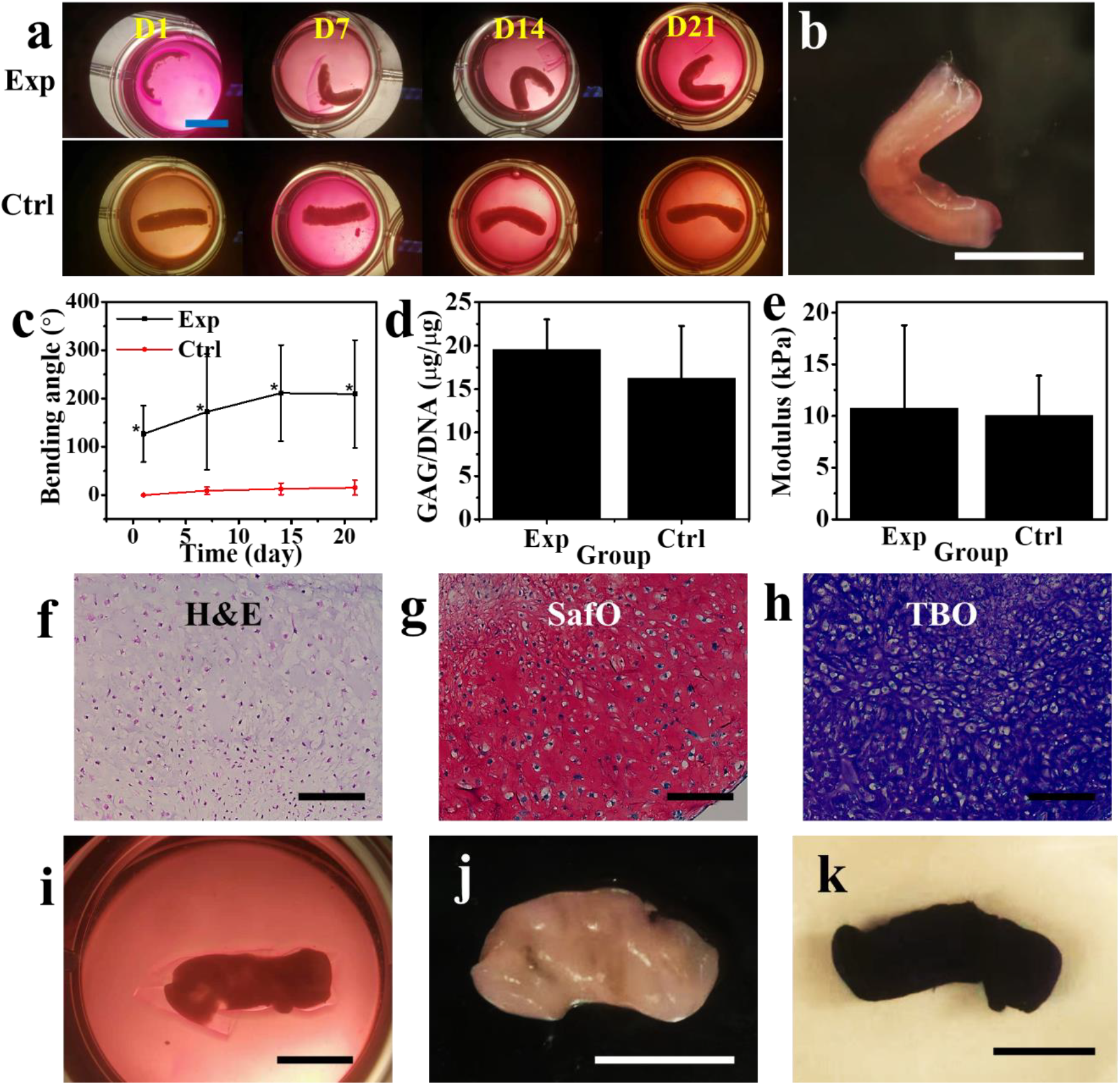
(a) Shapes of hMSC cell condensate-laden bilayer strips cultured in chondrogenic media at different times. Exp stands for the experimental group, Ctrl stands for the control group, and scale bar is 10 mm. The UV irradiation times for the gradient layer and the MG layer were set to 50 s and 20 s (50s/20s), respectively. (b) 4D engineered letter “C”-shaped cartilage-like tissue after 21 days of culture. Scale bar is 10 mm. (c) The change of bending angle as a function of the culturing time, \**p* < 0.05 compared to control group. (d) Biochemical analysis of GAG production (normalized to DNA content) at D21. (e) Young’s moduli of the ex vivo engineered cartilage-like tissues at D21. Histologic characterization of ex vivo 4D engineered cartilage-like tissues: (f) hematoxylin and eosin (H&E) staining, and GAG (g) Safranin O (SafO, pink/red) and (h) toluidine blue O (TBO, blue and purple) staining for GAG. Scale bars indicate 0.2 mm. Helix-shaped cartilage-like tissue obtained at day 21 (i) before and (j) after manual removal of outer hydrogel layer. Scale bars indicate 5 mm. The UV irradiation times for the gradient layer and the MG layer were set to 30 s and 20 s (30s/20s), respectively. (k) TBO stained helix-shaped cartilage-like tissue, scale bar is 5 mm.

As indicated earlier (Figure 2c and 2d), the shape of the bioconstruct can be tuned by adjusting the UV time for either the bottom layer or the upper layer. We demonstrated that increasing the UV time for the upper layer can also prolong the retention duration of the cell condensate layer before being released (Figure 5 and S15). To keep the same MG degradation profile and increase the deformability of the bilayer structure, we reduced the UV irradiation time from 50 s to 30 s to fabricate the bottom layer while maintaining a 20 s UV irradiation time for the upper layer, offering cell-strip laden bilayers (30s/20s UV crosslinking time) with greater curling, and yielding helix-shaped cartilage-like tissues after culturing for 21 days in chondrogenic media (Figure 6i and S21). Unlike the “C”-shaped cartilage-like tissues, the helix cartilage-like tissues were wrapped tightly with the gradient hydrogel (Movie S4) and did not separate spontaneously despite complete MG degradation. Since the formed cartilage-like tissues were very robust, they could be separated manually while maintaining their helix structures (Figure 6j and Movie S5). The whole construct also strongly stained with TBO due to the presence of a large amount of GAG (Figure 6k). The tunability illustrated herein highlights the versatility of this system to engineer cell condensation-based tissues with defined configurations on demand. Our results sufficiently demonstrated this 4D cell-condensate bioprinting strategy possesses high potential in the field of tissue engineering and may be expanded to other regenerative medicine related areas.

The 4D bioprinting technique opens a new avenue to engineer bioconstructs through a user-defined shape-morphing process, enabling dynamic 4D biofabrication at physiologically relevant timescales. Currently, in the 4D bioprinting field, research is still focused on the development of intelligent materials that undergo geometrical transformations under physiological conditions.^[44]^ Several reports to date have shown the feasibility of using cytocompatible polymers as bioinks to print high-resolution hydrogel constructs.^[30-35, 45]^ In these studies, cells were either seeded on the hydrogel surface or encapsulated inside the formed hydrogels. However, the seeding of cells on the hydrogel surface fails to replicate the 3D cellular microenvironment, while encapsulation of cells inside the hydrogels interferes with critical cell-to-cell interactions. Here we have proposed a novel 4D bioprinting strategy to address the challenges in the formation of geometrically complex scaffold-free 3D cell condensate constructs with defined configurations capable of undergoing temporally controllable and predefined architectural changes. Through a rational bilayer design using a gradient hydrogel as an actuation layer and a photocurable and biodegradable MG layer as a temporally controlled cell condensate-supporting layer, stable cell condensates with diverse and tunable conformational changes over time were obtained through programmable deformations. The cell condensate-derived tissues or other bioconstructs can be liberated after being cultured to a predetermined time point. As a proof-of-concept study, we demonstrated the fabrication of letter “*C*”-shaped and helix-shaped cartilage-like tissues. The unique features of this system on the whole lie in the (i) adjustable shape morphability, (ii) smooth and high-resolution MG printing and high-fidelity cell-only printability within the pre-printed MG layer, (iii) cytocompatible materials and processing, including 4D printing and shape transformations, and importantly (iv) controllable cell condensate formation, maturation, and release. This is the first time, to the authors’ best knowledge, that a 4D system has been implemented to achieve deformable cell condensations. We also observed very limited shape change caused by cell-contractile forces in the ex vivo 4D tissue engineering study. Given that cell-contractile forces can occur strongly within cell condensates and, as observed in this study, can be a natural trigger for impelling the 4D process, it might be of value to design a 4D cell-condensate bioprinting system that involves the cell-contractile force as a collaborative, primary or even the sole stimulus.

## 4. Conclusions

In this work, we have constructed a unique 4D bioprinting system to produce specifically shaped cell condensates for applications such as developmental mimicry and tissue engineering. The establishment of this strategy may shed valuable insights into the future development of 4D bioprinting associated with scaffold-free tissue regeneration. This system addresses the issues associated with the use of scaffold-based 4D tissue engineering approaches. Our strategy is effective in enabling construction of scaffold-free cell condensates with diverse geometries that can be designed to undergo preprogrammed shape changes over time, offering a versatile platform for customized biofabrication.

## Experimental Section/Methods

Please refer to the supporting information for the experimental section.

## Supporting Information

Supporting Information is available from the Wiley Online Library or from the author.

## Acknowledgements

The authors gratefully acknowledge funding from the National Institutes of Health’s National Institute of Arthritis and Musculoskeletal and Skin Diseases (R01AR074948) and National Institute of Biomedical Imaging and Bioengineering (R01EB023907). The contents of this publication are solely the responsibility of the authors and do not necessarily represent the official views of the National Institutes of Health.

Received: ((will be filled in by the editorial staff))

Revised: ((will be filled in by the editorial staff))

Published online: ((will be filled in by the editorial staff))

## Notes

### Competing Interest Statement

The authors have declared no competing interest.

