## Supporting Information for "4D Cell-Condensate Bioprinting"

#### 1. Experimental:

#### 1.1 Chemicals, instruments, and general methods

Unless specified, all solvents and reagents were used without further purification. Sodium alginate (**AL**, Protanal LF120M, 157 Pa·s and 251 Pa·s) was a generous gift from FMC Biopolymer. Bovine skin derived gelatin (type B), photoinitiator (2-Hydroxy-4'-(2-hydroxyethoxy)-2-methylpropiophenone, PI), 4'-hydroxy-3'-methylacetophenone (HMAP), fluorescein diacetate (FDA), ethidium bromide (EB), Dulbecco's Modified Eagle Medium-Low Glucose (DMEM-LG), and fetal bovine serum (FBS) were purchased from Sigma. $ITS^+$ Premix and penicillin/streptomycin (P/S) were purchased from Corning Inc. (Corning, NY). Sodium pyruvate was purchased from HyClone Laboratories. Non-essential amino acid solution was purchased from Lonza Group (Basel, Switzerland). Ascorbic acid-2-phosphate was purchased from Wako Chemicals USA Inc. (Richmond, VA). Fibroblast growth factor-2 (FGF-2) was purchased from R&D Systems (Minneapolis, MN). Transforming growth factor β1 (TGF-β1) was purchased from PeproTech (Rocky Hill, NJ). *N*-(2-aminoethyl) methacrylate hydrochloride (AEMA) and methacryloxyethyl thiocarbamoyl rhodamine B (RhB) were purchased from Polysciences Inc., and other common chemicals, such as sodium peroxide, methacrylic anhydride, etc., were purchased from Fisher Scientific. $^1H$ NMR spectra were obtained on a 400 MHz Bruker AVIII HD NMR spectrometer equipped with a 5 mm SmartProbe™ at 25 ºC using deuterium oxide ($D_2O$) as a solvent and calibrated using (trimethylsilyl)propionic acid-$d_4$ sodium salt (0.05 w/v %) as an internal reference. DMEM-LG containing 0.05% PI (w/w) was used to dissolve the oxidized methacrylate alginate (OMA) and methacrylate gelatin (GelMA). Cell growth media (GM) consisted of DMEM-LG with10% FBS and 1% P/S, and chondrogenic media consisted of DMEM-LG with 1% $ITS^+$ Premix, 100 nM dexamethasone, 1 mM sodium pyruvate, 100 µM non-essential amino acids, 37.5 µg/mL ascorbic acid-2-phosphate and 1% P/S supplemented with 10 ng/mL TGF-β1.

Hydrogel images were visualized using a Nikon SMZ-10 Trinocular Stereomicroscope equipped with a digital camera. A microplate reader (Molecular Devices iD5) was used to read data from the microplates. A UV device (EXFO OmnicureR S1000-1B, Lumen Dynamics Group) with an intensity of 12 mW/cm$^2$ was used for photocrosslinking. All quantitative data was expressed as mean ± standard deviation. Statistical analysis was performed with one-way analysis of variance (ANOVA) with Tukey honestly significant difference post hoc tests using Origin software (OriginLab Corporation). A value of $p < 0.05$ was considered statistically significant.

**1.2 Synthesis of OMAs and GelMA**

OMAs with a theoretical 5% oxidation degree and varying theoretical methacrylation degrees (20%, and 45%) were synthesized according to a similar method as described in the literature.[S1-2] The O5M20A (5% theoretical oxidation and 20% theoretical methacrylation) was synthesized with the following procedure: 10 g of sodium alginate (251 Pa·s) was dissolved in 900 mL of di$H_2O$ overnight, and 0.54 g of sodium periodate ($NaIO_4$) in 100 mL of di$H_2O$ was rapidly added to the alginate solution under stirring in the dark at room temperature (RT). After reaction for 24 h, 19.52 g of 2-ethanesulfonic acid (MES) and 17.53 g of sodium chloride (NaCl) were added, and the pH was adjusted to 6.5 with 5 N sodium hydroxide (NaOH). Then 1.18 g of *N*-hydroxysuccinimide (NHS) and 3.89 g of 1-ethyl-3-(3-dimethylaminopropyl)carbodiimide hydrochloride (EDC·HCl) were sequentially added to the mixture. After 10 min, 1.69 g of AEMA was added slowly. The solution was wrapped with aluminum foil to protect it from light and left to react for 24 h at RT. The mixture was then poured into 2 L of chilled acetone to precipitate out the crude OMA solid, which was further purified by dialysis against di$H_2O$ over 3 days (MWCO 3.5 kDa, Spectrum Laboratories Inc.). The dialyzed alginate solution was collected, treated with activated charcoal (0.5 mg/100 mL, 50-200 mesh, Fisher) for 30 min, filtered through a 0.22 μm filter and frozen at -80 ºC overnight. The final O5M20A was obtained as white cotton-like solid through lyophilization for at least 10 days. O5M45A (5% theoretical oxidation and 45% theoretical methacrylation) was synthesized through the same procedure according to the reported literature.[S3] The actual methacrylation of O5M20A and O5M45A was determined to be 5.7% and 16.2% from $^1$H NMR data according to the method described in the literature. [S4] Note that the actual oxidations were not provided due to the overlap of the proton peak assigned to the CHO group (~5.4 ppm) with the polymer proton peak (broad peak located at ~5.1 ppm). The

GelMA was the same material that was used in our previous work.[S3] The $^{1}H$ NMR spectra for newly synthesized OMAs are shown in Figure S22 and S23.

#### 1.3 Microgel (MG) preparation

MGs was prepared using a modified procedure from the reported literature. [S5] To make the stock MGs, O5M20A (1.2 g) was dissolved in deionized water (di$H_2O$, 60 mL) and then was slowly dispensed into a gelling bath containing an aqueous solution of $CaCl_2$ (600 mL, 0.2 M) under fast stirring with a magnetic stir bar. After fully ionically crosslinking overnight, the resultant hydrogel beads were collected, washed once with 40 mL of 70% ethanol (EtOH)/water ($H_2O$), and then blended twice using a household blender (Osterizer MFG, at "pulse" speed) for 2 min with 120 mL of 70% EtOH/$H_2O$. Then, the MGs were obtained and loaded into 50 mL conical tubes and centrifuged at 4200 rpm for 5 min and stored at 4 °C for future use.

To recover the MGs for use, the as-prepared MGs (5 mL) in 70% EtOH/$H_2O$ were washed 3 times by replacing the previous media with 25 mL of 0.05% (*w*/*w*) PI-contained di$H_2O$ and then vortexed (Fisher STD Vortex Mixer, Fisher Scientific, 10× speed) for 2 min every time and subsequently washed 1 time by replacing the previous media with 25 mL of 0.05% (*w*/*w*) PI-containing DMEM-LG and vortexed (10× speed) for 10 s. This recovered MGs were used for further experiments.

#### 1.4 Gradient and non-gradient hydrogel preparation

Gradient hydrogels were used as the actuation layer to drive shape morphing. The preparation was according to a similar procedure described in the reported literature.[3] Briefly, a mixed solution of polymer [OMA 6% w/w, OMA (6% w/w)/GelMA (6% w/w)], PI (0.05% w/w), and UV absorber [RhB (0.01% w/v)/HMAP (0.02%)] in DMEM-LG was placed between two quartz plates with 0.6 mm spacers and subsequently photocrosslinked with UV light (EXFO OmnicureR S1000, Lumen Dynamics Group) at 12 mW/cm$^2$ for a certain time (20 s ~ 40 s) to form a O5M45A hydrogel disc with a gradient in crosslinking density throughout the thickness, which was denoted as O5M45A(g) or O5M45A/GelMA(g). Non-gradient hydrogels were used for comparison and were fabricated similarly without using a UV absorber and were denoted as O5M45A or O5M45A/GelMA. The gradient/non-gradient hydrogels discs were cut into hydrogel sheets with dimensions of 25 mm × 25 mm as a substrate for the MG printing (described later).

#### 1.5 Microgel (MG) printing

The MG printing was performed using a 3D printer (PrintrBot Simple Metal 3D Printer, Printrbot) modified with a syringe-based extruder. More information about this printer can be found in the literature.[S5] The STL files for the bioink printing were generated from www.tinkercad.com.

The MGs were loaded into a 1 mL glass syringe (Hamilton, Reno, NV), which was connected to a 22-gauge (22G) stainless-steel needle (McMaster-Carr) and mounted into the syringe pump extruder on the 3D printer. The above O5M45A(g) hydrogel sheet was flipped and placed on a quartz plate with low-crosslinking side attaching the quartz plate surface. The tip of the needle was positioned at the center and near the surface of the O5M45A or O5M45A(g) hydrogel sheet, and the print instructions were sent to the printer using the host software (Cura Software, Ultimaker), which is an open-source 3D printer host software. The MGs were printed under 4 mm/s printing speed with 100% infilling density. After printing, the obtained constructs were used for further cell-only printing (described later) or immediately photocured under UV irradiation. Then, the cell-free constructs were cut into specific shapes (*e.g.*, strip, sheet, and disc). These hydrogels were then carefully transferred into a tissue culture plate for culturing to monitor the shape change or/and allow cell differentiation. The hydrogels were imaged, and the bending angles were quantified according to the previous literature.[S6]

For printing MGs for swelling and degradation studies, Young's modulus testing, rheological testing, and the cell printability study, MGs were printed according to a similar method described above. The MGs were directly printed on the quartz plate instead of on the surface of a gradient hydrogel.

#### 1.6 Bilayer hydrogel fabrication

Gradient/non-gradient hydrogels as substrates were fabricated as above (Section 1.4) and MGs were printed on the surface of the hydrogel substrates as above (Section 1.5). The bilayer hydrogels were then photocrosslinked for a certain time. To distinguish the UV time for the bilayer fabrication, the total UV time was denoted as gradient layer time/MG layer time. For example, if the UV time for the gradient hydrogel layer and the MG layer is set to 30 s and 20 s, respectively, the overall UV time for the bilayer fabrication is denoted as 30s/20s. The bilayer is denoted as bottom hydrogel_upper hydrogel. For example, O5M45A/GelMA(g)_O5M20A MG is a bilayer consisting of O5M45A/GelMA(g) (bottom layer) and O5M20A MG (upper layer).

### 1.7 Swelling and degradation tests

Gradient/non-gradient hydrogels and UV crosslinked MGs were used for swelling and degradation tests. As described above, gradient/non-gradient hydrogels were fabricated with a specific UV crosslinking time of 60 s, and MGs were printed into cuboids with dimensions of 8 × 6 × 3 $mm^3$ and then UV crosslinked for 20 s or 30 s. The hydrogel samples were frozen for 4 h at -80 ºC and lyophilized for 2 days. The masses of the dried gels were measured as initial weights (*W*i). The dried hydrogels were rehydrated by culturing in 5 mL of GM at 37 °C, and the media was changed every 3 days. At predetermined time point, the hydrogels were collected, and the swollen weights (*Ws*) were measured. The swelling ratio was calculated with the following equation: *Ws*/*Wi* (N = 3). For the degradation test, the rehydrated hydrogels were collected at predetermined time points over 28 days and dried by lyophilization to obtain dried mass ($W_d$). Mass loss was quantified as ($W_i$-$W_d$)/$W_i$ × 100% (N = 3) for each condition per time point.

### 1.8 Young's modulus measurement

The elastic moduli of the gradient/non-gradient hydrogels and UV crosslinked MGs were determined by performing uniaxial, unconfined constant strain rate compression testing at RT using a constant crosshead speed of 0.8%/sec on a mechanical testing machine (225lbs Actuator, TestResources, MN, USA) equipped with a 5 N load cell. To obtain the Young's modulus of the engineered cartilage-like tissue (described later), the tissues were punched into a cylinder ($d$ = 2 mm, $h$ = 1.2 mm) using a biopsy punch. The compression tests were performed with the same protocol as above except a 0.5%/sec crosshead speed was used. The Young's modulus of each sample was determined using the first non-zero slope of the linear region of the stress-strain curve within 0~10% strain (N = 3).

### 1.9 Rheological testing

Dynamic rheological examination of the uncrosslinked and photocrosslinked MGs was performed to measure the hydrogel storage and loss moduli and viscosity and evaluate the shear-thinning, shear yielding, and self-healing properties with a Kinexus ultra+ Highest specification rheometer (Malvern Panalytical). In oscillatory mode, a parallel plate geometry (8 mm diameter) measuring system was employed, and the gap was set to 1 mm. After each hydrogel was placed between the plates, all the tests were carried out at RT. Oscillatory frequency sweep (0.1~100 Hz at 1 % strain) tests were performed to measure storage moduli

(G'), loss moduli (G"), and viscosity. Oscillatory strain sweep (0.01~100 % strain at 1 Hz) tests were performed to examine the shear-thinning characteristics of the MGs and to determine the shear-yielding points at which the MGs behave fluid-like. To investigate self-healing properties, cyclic deformation tests were performed at 100% strain with recovery at 1% strain, each for 1 min at 1 Hz.

**2.0 Interfacial adhesion testing**

The tensile testing was performed according to the reported literature using a mechanical testing machine (225lbs Actuator, TestResources, MN, USA) equipped with a 25 N load cell to evaluate the interfacial adhesive strength of the bilayer hydrogels (O5M45A/GelMA(g)_O5M20A).[6] Briefly, the hydrogel samples with an interfacial cross-sectional area of 5 × 1 $mm^2$ were attached to two hard paper backings using cyanoacrylate glue (Krazy Glue®, Elmer's Products Inc., Columbus, OH). The hard paper backings were then attached firmly with common commercial transparent tape to a "plastic loading platen", which was attached to the "load cell", and to a "sample cup", which was fixed on the bottom platform of the mechanical testing machine with a 5 mm gap. The adhesion strength was determined by performing constant strain rate (0.6%/sec) tensile tests at RT.

**2.1 Cell expansion**

Human mesenchymal stem cells (hMSC) were isolated according to the literature.[S7] hMSC cells were expanded in GM supplemented with 10 ng/mL FGF-2 and HeLa cells were expanded in GM. The incubation was performed in a humidified incubator at 37 °C and 5% $CO_2$ with media changes every 2 or 3 days. The cells were harvested as cell-only bioinks according to the protocol described in the literature[5] for printing when they reached ~80% confluence.

**2.2 Cell-only bioprinting**

HeLa and hMSC cells as bioinks were loaded into 3 mL of luer lock syringes (Becton, Dickinson and Company, NJ), connected to a 25G stainless steel needles (McMaster-Carr) and mounted into the BIO X 3D printer (CELLINK, MA). Pre-printed MGs described above (Section 1.5) were used as the supporting batch for 3D cell-only printing. The printing parameters were set to 95% infilling density, 2 mm/s printing speed, and 0.8 μL/s (HeLa cells) or 1.0 μL/s (hMSC cells) extrusion rate. After 3D printing of the cell-only bioinks, cell-laden O5M20A MGs were stabilized by photocrosslinking under UV for a specified time. The photocrosslinked cell-laden bioconstructs were transferred into a 4-well tissue culture plate for culturing the strip-shaped hydrogels with 4 mL of GM or a 6-well tissue culture plate for culturing bioconstructs with other shapes with 10 mL of GM to record shape change and assess cell viability through a live/dead staining assay (described later). The media was changed every day.

#### 2.3 Live/dead staining

The viability of cells was assessed using live/dead staining comprised of FDA and EB. The staining solution was freshly prepared by mixing 1 mL of FDA solution (1.5 mg/mL in DMSO) and 0.5 mL of EB solution (1 mg/mL in PBS) with 0.3 mL PBS (pH 8). At predetermined time points, 20 μL of staining solution per 1 mL of culture media was added into each well and incubated for 5 min at RT. Fluorescence images of the samples were taken using a Nikon Eclipse TE300 fluorescence microscope (Nikon, Japan) equipped with a 14MP Aptina Color CMOS digital camera (AmScope, CA).

#### 2.4 Ex vivo cartilage-like tissue engineering, biochemical quantification, and histological staining.

The hMSCs at passage 4 (P4) were used for the chondrogenesis study. The cells were harvested for cell-only bioprinting when they reached ~80% confluence. The cell-laden bilayer hydrogel strips were fabricated as above (Section 1.4 −1.6) and cultured in chondrogenic media. The UV application time for the letter “C”- and helix-shaped bioconstruct formation was set to 50s/20s and 30s/20s, respectively. For cartilage-like tissue formation, the hMSC-laden hydrogels were cultured in 4 mL of chondrogenic media in a humidified incubator at 37 °C with 5 % $CO_2$ over a course of 21 days, and 2 mL of media was changed every day. The cartilage-like tissues were obtained at day 21 and cut into small pieces for biochemical analysis, Young’s modulus testing (Section 1.8), and histological staining.

For biochemical analysis, the small tissue pieces were digested in 0.5 mL of papain solution (Sigma) at 65 ºC for 24 hours[8] and centrifuged for 10 min at 15,000 rpm, and then the supernatants were collected for DNA and glycosaminoglycan (GAG) quantifications (N = 3).

Per the manufacturer’s instructions, a Picogreen assay kit (Invitrogen) was used to quantify the DNA content in the supernatant. Fluorescence intensity of the dye-conjugated

DNA solution was measured using a microplate reader with an excitation of 480 nm and emission of 520 nm.

The GAG content was quantified using a DMMB (1,9-dimethylmethylene blue) assay.[S5] 40 μL of supernatant from the digested samples was transferred into 96-well plate, to which 125 μL of DMMB solution was then added. Absorbance at 595 nm was recorded on a microplate reader. GAG content was normalized to DNA content.

The small cartilage-like tissues were fixed with 10% neutral buffered formalin (NBF) overnight at 4 °C, dehydrated, and embedded in paraffin. Briefly, tissue samples were cut into 5 μm thick sections using a Leica RM2255 rotary microtome (Leica Microsystem Ltd., Milton Keynes, UK). Slide sections were then deparaffinized and stained with hematoxylin and eosin (H&E) stains to observe gross cell and tissue morphology,[S8] and safranin O (SafO) with a Fast Green counterstain[S9] and toluidine blue O (TBO) for glycosaminoglycan (GAG) indication.[S10] Stained samples were imaged using a fluorescence microscope (Nikon Eclipse TE300, Japan) under bright field.

TBO staining was also used to stain the entire helix-shaped cartilage-like tissue. The whole tissue was collected and stained with Toluidine blue O for 30 min and washed with PBS 3 times to remove the unbound stain.[S5]

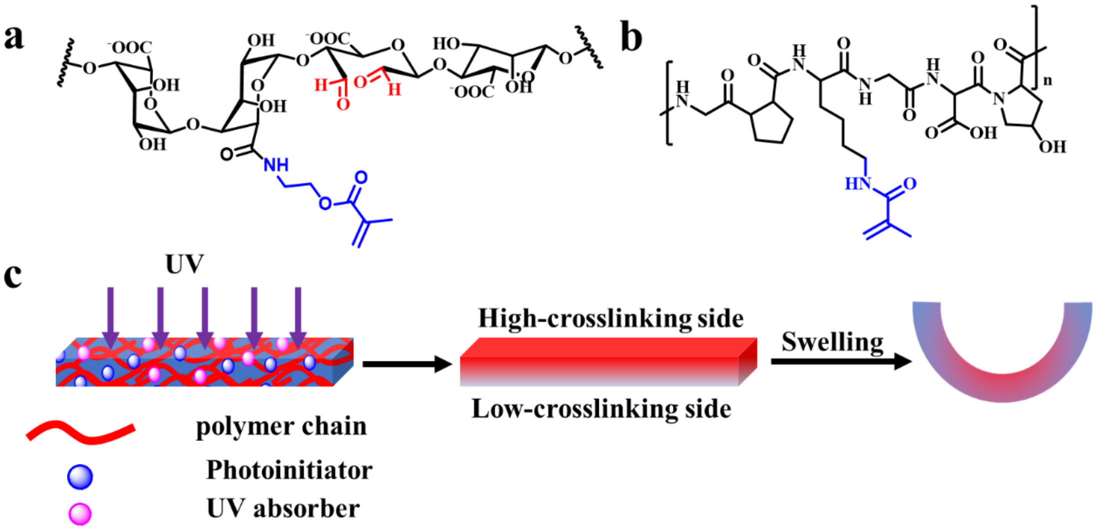


Figure S1. Chemical structure of (a) OMA and (b) GelMA. (c) Schematics showing the gradient hydrogel formation and its bending after swelling in media.

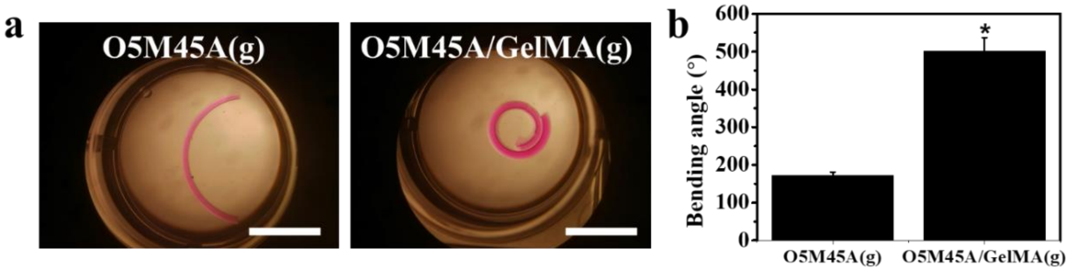


Figure S2. (a) Representative images showing curled O5M45A(g) (left) and O5M45A/GelMA(g) (right) hydrogel strips after culture in PBS at RT for 30 s. (b) The corresponding bending angles of the O5M45A(g) and O5M45A/GelMA(g) hydrogel strips. $*p < 0.05$, scale bars indicate 10 mm.

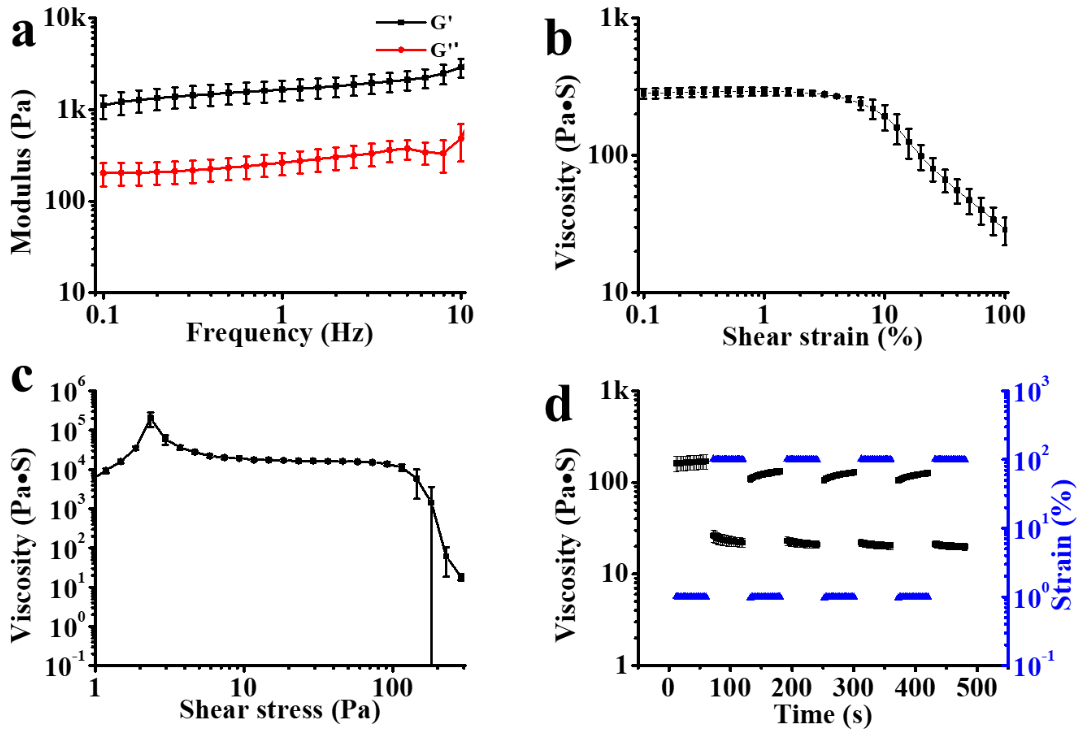


Figure S3. (a) The change of storage (G’) and loss (G’’) moduli of uncrosslinked O5M20A MG with increasing sweep frequency. The change of viscosity with increase of (b) shear strain and (c) shear stress. (d) Reversible viscosity under alternating shear strain between 1% and 100%.

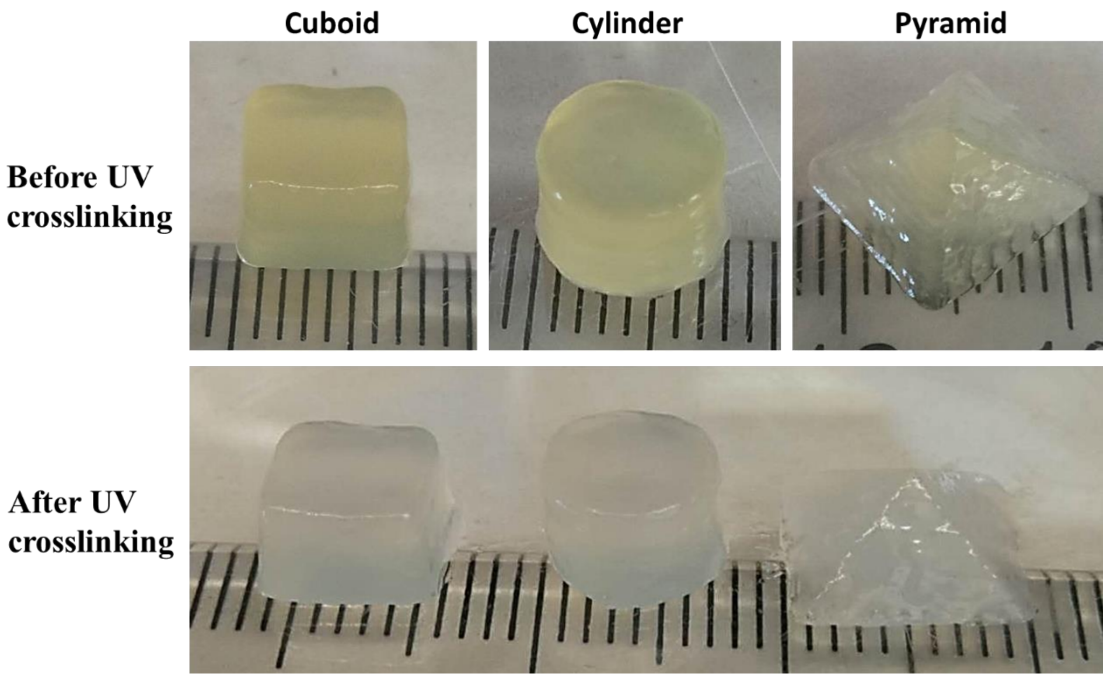


Figure S4. 3D printed O5M20A MG constructs (a) before UV crosslinking and (b) after UV crosslinking. Cuboid: length × width × height = 7 mm × 7 mm × 6 mm. Cylinder: diameter × height = 8 mm × 6 mm. Pyramid: bottom length × height: 10 mm × 4 mm.

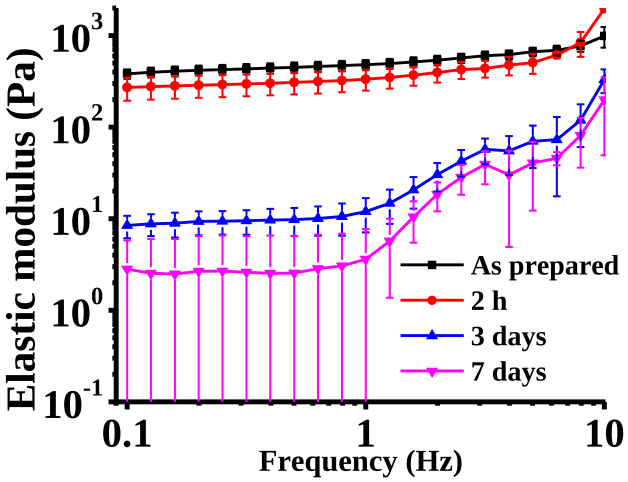


Figure S5. The change of elastic modulus of the crosslinked O5M20A MGs with culture time in the GM under a cell-culture environment.

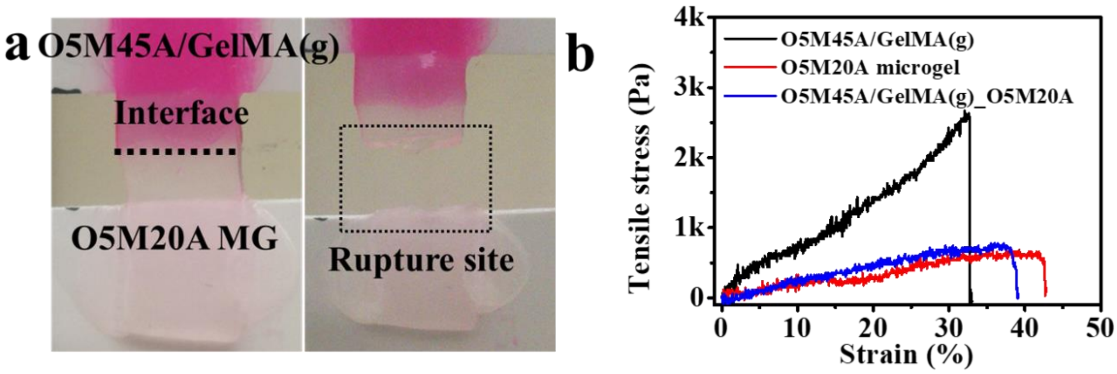


Figure S6. (a) Images of O5M45A/GelMA(g)_O5M20A MG bilayer before (left) and after (right) rupture in a tensile test. The interfacial adhesion strength is stronger than the strength of the O5M20A MG, and thus it ruptured on the O5M20A MG side instead of at the interface. (b) Representative tensile stress-strain curves of the O5M45A/GelMA(g) hydrogel, the O5M20A MG hydrogel, and the O5M45A/GelMA(g)_O5M20A MG bilayer hydrogel.

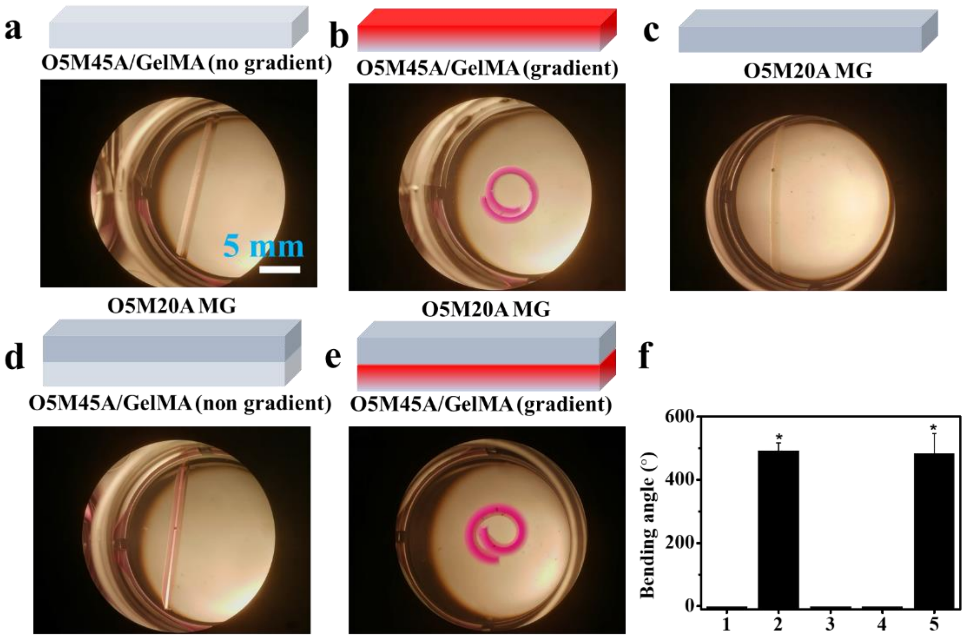


Figure S8. Representative images showing the shape of (a) O5M45A/GelMA, (b) O5M45A/GelMA(g), (c) O5M20A MG, and (d) O5M45A/GelMA_O5M20A MG, (e) O5M45A/GelMA(g)_O5M20A MG. (f) The bending angle of the respective single-layer and bilayer hydrogels, 1: O5M45A/GelMA; 2: O5M45A/GelMA(g); 3: O5M25A MG; 4: O5M45A/GeMA_O5M20A MG; 5: O5M45A/GelMA(g)_O5M20A MG. *$p < 0.05$ compared to groups lacking a symbol.

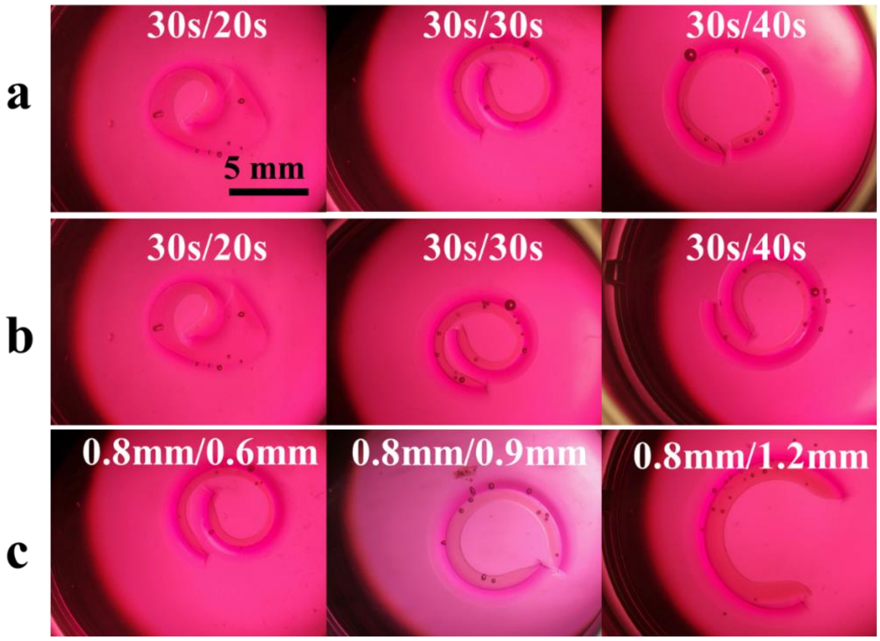


Figure S9. Representative photomicrographs of hydrogel strips fabricated under different conditions: varying the UV application time for (a) the upper layer (O5M20A MG layer) and (b) the bottom layer (O5M45A/GelMA gradient layer) and (c) varying the MG layer thickness (bottom layer/top layer). The hydrogel strips were cultured in DMEM-LG at room temperature for 30 min to reach equilibrated swelling. The hydrogel strip thickness for UV time variation study was set to bottom/top 0.8/0.5 (mm/mm) and the UV time for the MG layer thickness variation study was set to bottom/top 30s/30s.

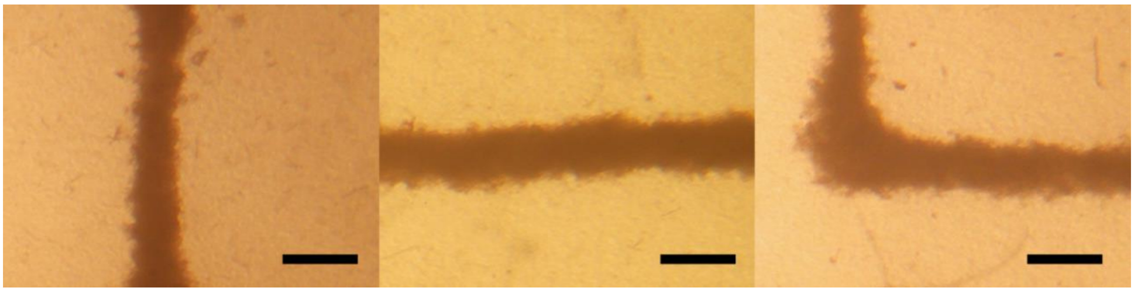

Figure S10. Photomicrographs of the printed cell filaments inside O5M20A MG after 20 s UV crosslinking. Scale bars indicate 0.5 mm.

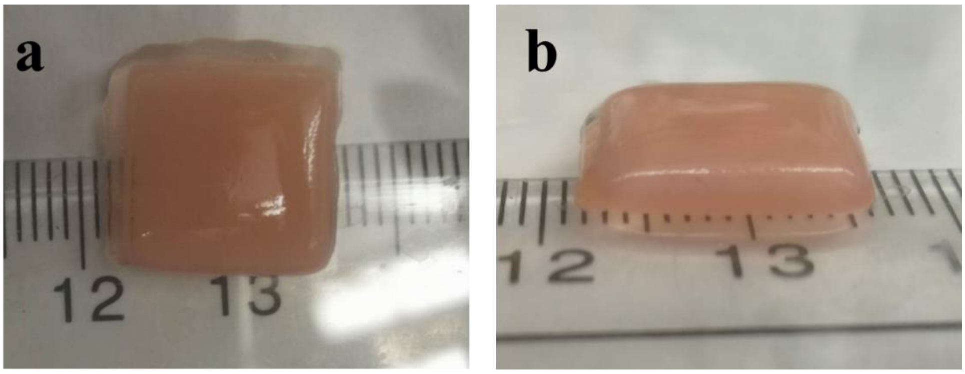


Figure S11. Photographs of as-printed HeLa cell sheet inside the O5M20A MG before UV crosslinking: (a) top view, (b) front view.

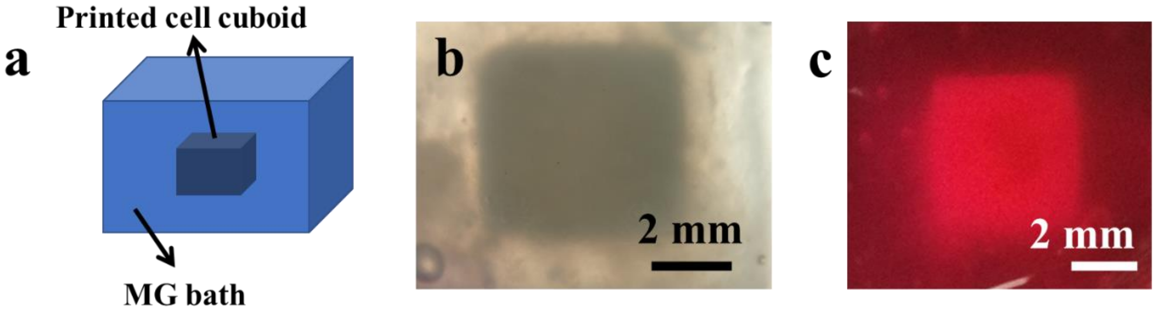


Figure S12. (a) Illustrated image and (b) photomicrograph of a printed HeLa cell cuboid inside the photocrosslinked O5M20A MG. (c) Photomicrograph of the cell tetrahedron cultured in the cell growth media after UV crosslinking. Cell tetrahedron size: 5 mm × 5 mm × 4 mm.

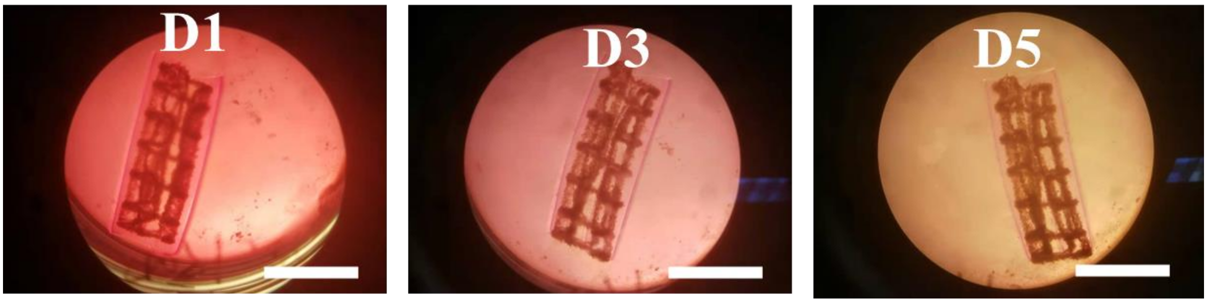


Figure S13. Photomicrographs of the HeLa cell-net infilled bilayer hydrogel sheet at D1, D3, and D5. The bioconstruct was cultured in 10 mL of cell growth media. Scale bars indicate 10 mm. UV crosslinking time was set to 40s/20s.

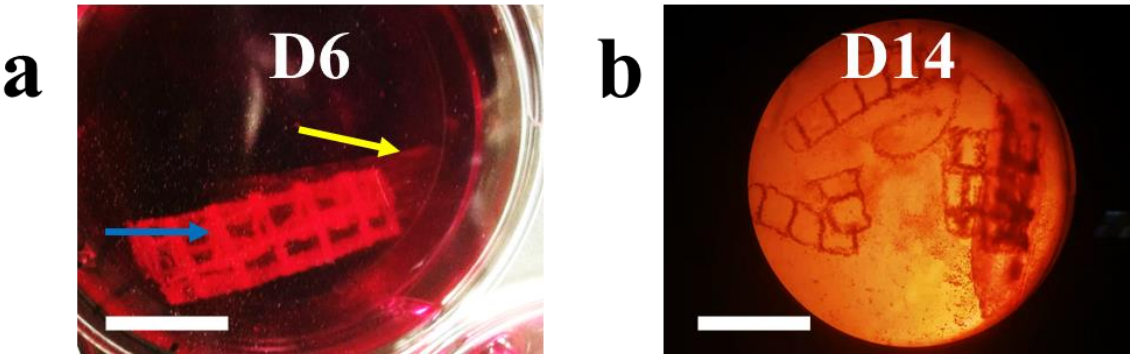


Figure S14. (a) Photograph of a HeLa cell-net infilled bilayer hydrogel sheet at D6, yellow and blue arrows show the separated gradient hydrogel layer and the cell condensate-laden MG layer, respectively. (b) Photomicrograph of a cell-net infilled bilayer hydrogel sheet at D14. The bioconstruct was cultured in cell growth media. Scale bars indicate 10 mm. UV crosslinking time was set to bottom/top 40s/20s.

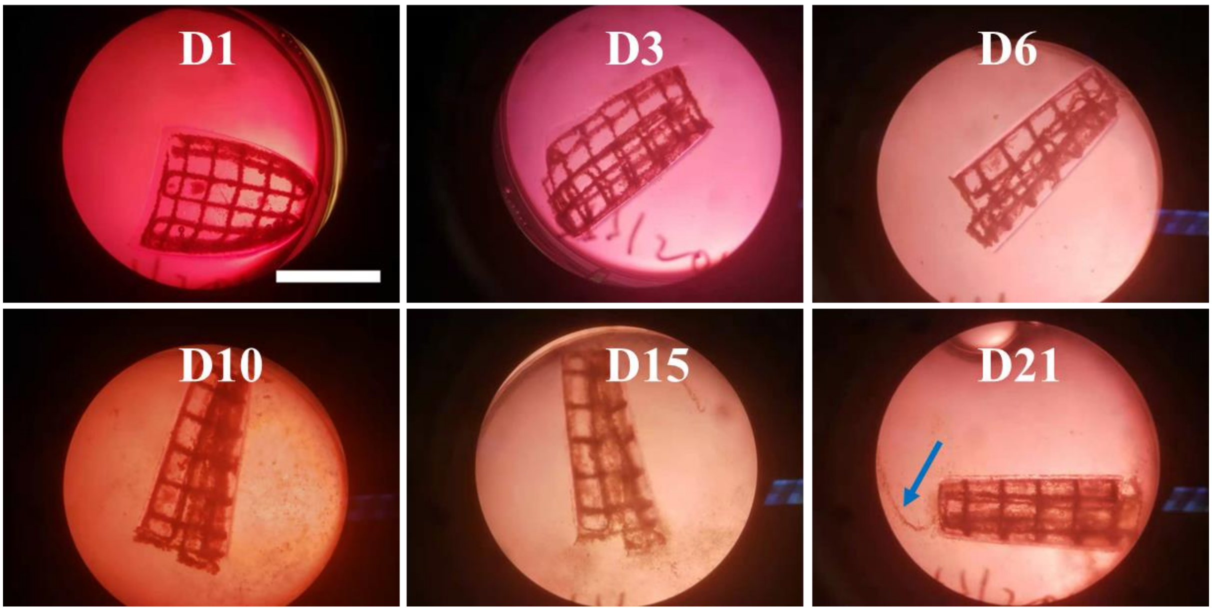


Figure S15. Photomicrographs showing the morphological changes of a HeLa cell-net filled bilayer hydrogel sheet over 21 days of culture in cell grow media. Arrow at D21 shows a “liberated” cell filament. Scale bar is 5 mm. UV crosslinking time was set to bottom/top 40s/30s.

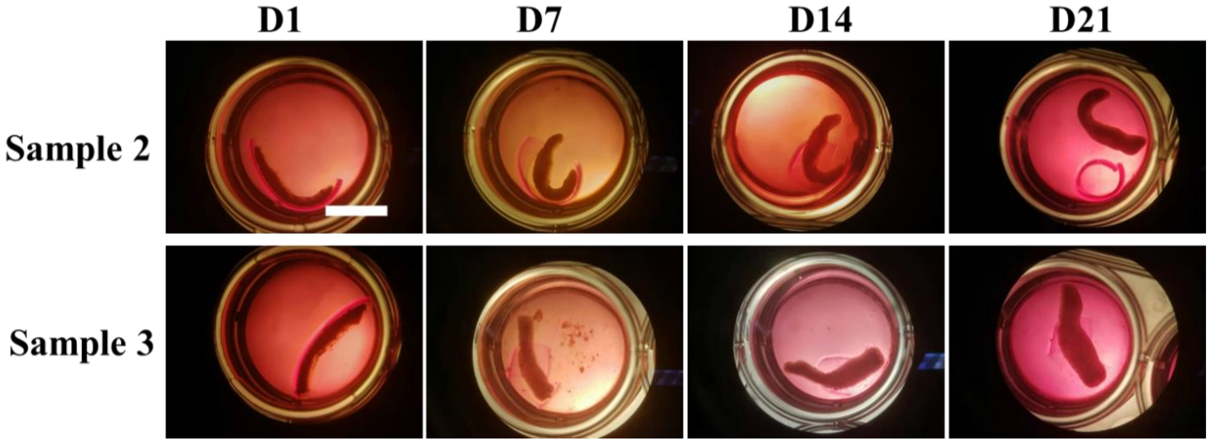


Figure S16. Shape changes of another two hMSC cell-laden bilayer strips over 21 days of culture and cell condensation formation in chondrogenic media. Scale bar is 10 mm. UV crosslinking time was set to bottom/top 50s/20s.

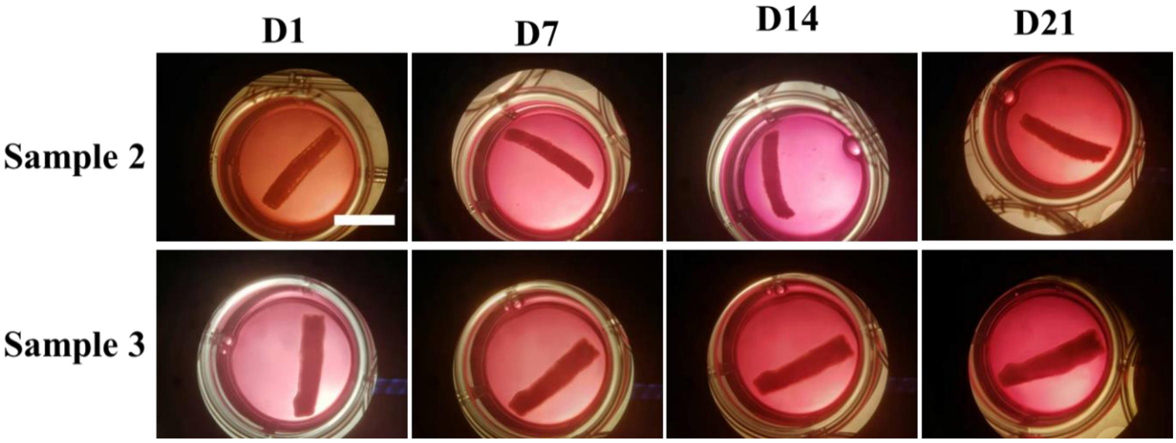


Figure S17. Shape changes of another two hMSC cell-laden single-layer O5M20A MG strips over 21 days of culture and condensation formation in chondrogenic media. 20 s UV crosslinking time. Scale bar is 10 mm.

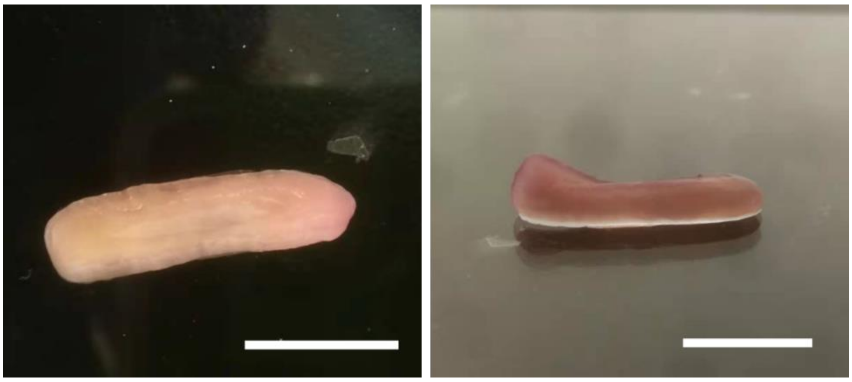

Figure S18. Photographs of a near straight line-shaped cartilage-like tissue obtained from the control group (hMSCs in photocrosslinked O5M20A MGs after 21 days culture in chondrogenic media): top view (left) and front view (right). Scale bars indicate 10 mm.

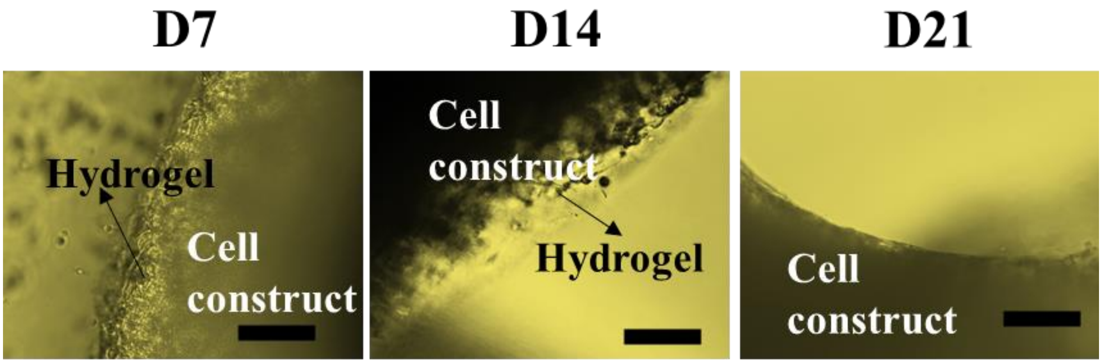


Figure S19. Photomicrographs showing the degradation of the photocrosslinked supporting O5M20A MGs during the 21 days of culture. Scale bars indicate 0.2 mm.

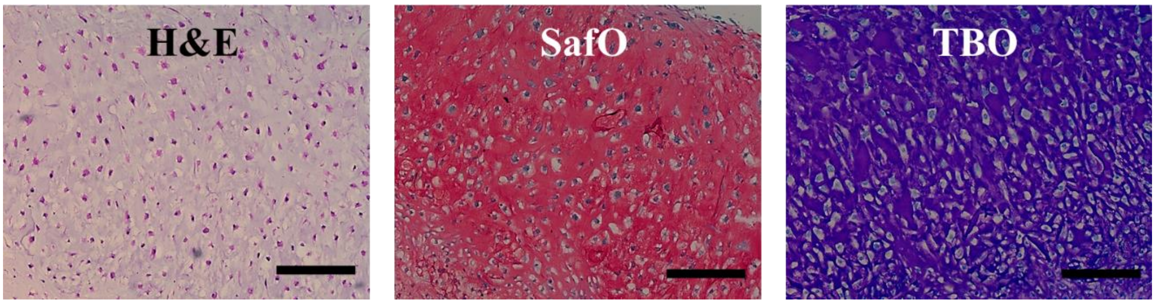


Figure S20. Histological characterization of cartilage like tissue obtained from the control group (hMSCs in photocrosslinked O5M20A MG after 21 days culture in chondrogenic media): hematoxylin and eosin (H&E) staining for gross cell and tissue morphology, safranin O (SafO, red/pink) and toluidine blue O (TBO, blue and purple) staining for GAG. Scale bars indicate 200 μm.

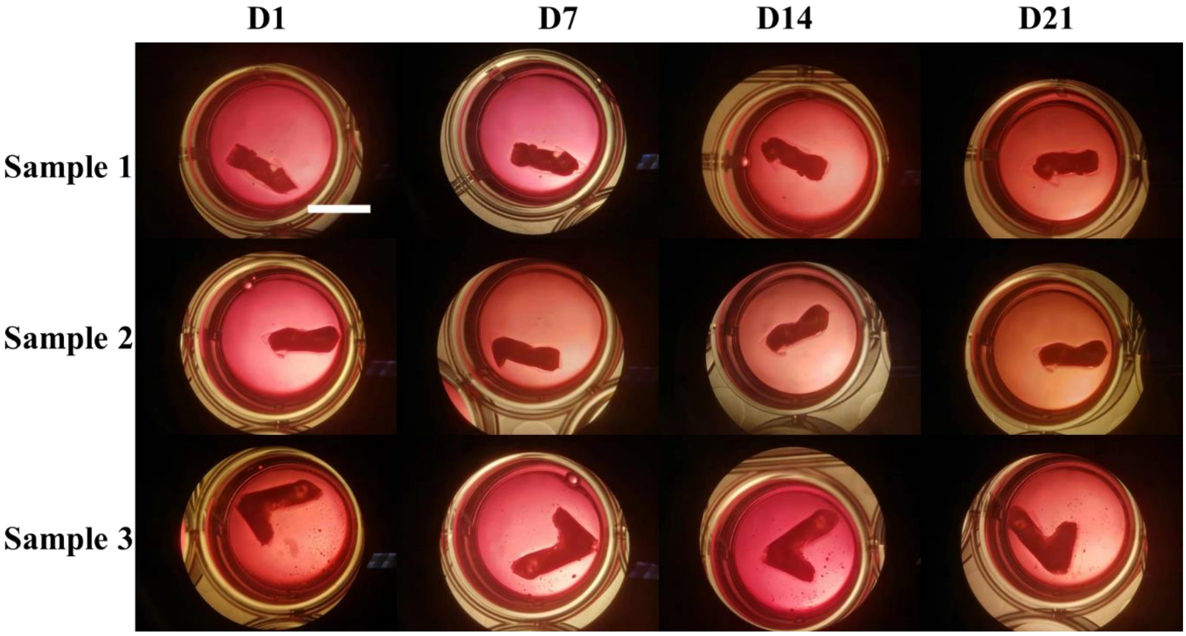


Figure S21. Morphologies of helix-shaped cell-laden bilayer hydrogels over 21 days of culture and cell condensation formation in chondrogenic media. UV crosslinking time was set to bottom/top 30s/20s. Scale bar is 10 mm.

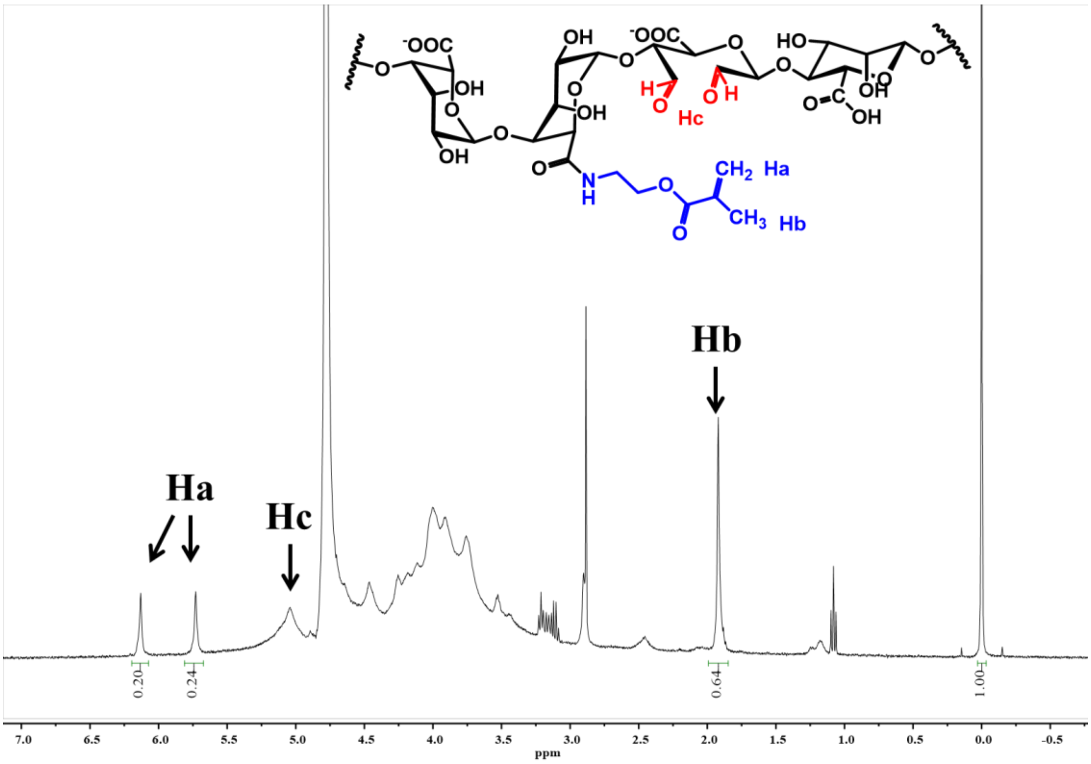


Figure S22. $^{1}H$ NMR spectrum of O5M20A ($D_2O$, 2 w/v %).

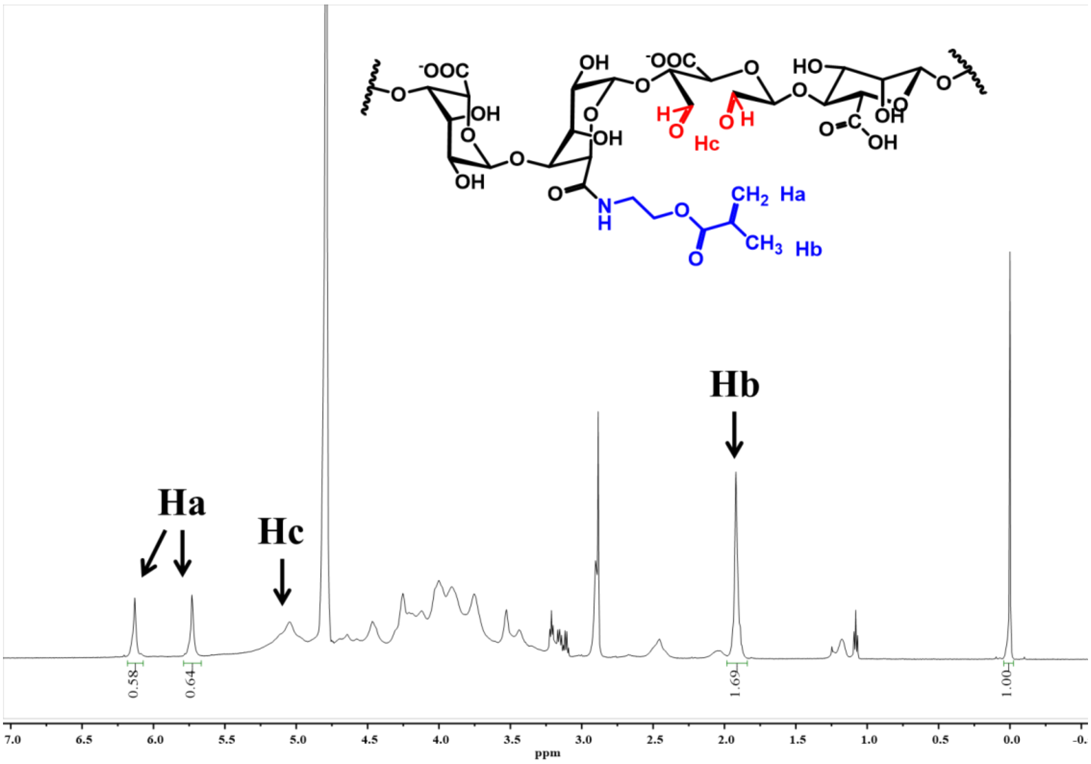


Figure S23. $^{1}H$ NMR spectrum of O5M45A ($D_2O$, 2 w/v %).

(S8) Dikina, A. D.; Alt, D. S.; Herberg, S.; McMillan, A.; Strobel, H. A.; Zheng, Z.; Cao, M.; Lai, B. P.; Jeon, O.; Petsinger, V. I. A modular strategy to engineer complex tissues and organs. *Advanced science* **2018,** *5* (5), 1700402.

(S9) Dikina, A. D.; Strobel, H. A.; Lai, B. P.; Rolle, M. W.; Alsberg, E. Engineered cartilaginous tubes for tracheal tissue replacement via self-assembly and fusion of human mesenchymal stem cell constructs. *Biomaterials* **2015,** *52*, 452-462.

(S10) Wu, H.; Shen, L.; Zhu, Z.; Luo, X.; Zhai, Y.; Hua, X.; Zhao, S.; Cen, L.; Zhang, Z. A cell-free therapy for articular cartilage repair based on synergistic delivery of SDF-1 & KGN with HA injectable scaffold. *Chemical Engineering Journal* **2020,** *393*, 124649.
